# Sex-dependent effects of early life stress on network and behavioral states

**DOI:** 10.1101/2024.05.10.593547

**Authors:** Garrett Scarpa, Pantelis Antonoudiou, Grant Weiss, Bradly Stone, Jamie L. Maguire

## Abstract

**Background:** Adverse childhood experiences (ACEs) are associated with numerous detriments in health, including increased vulnerability to psychiatric illnesses. Early life stress (ELS) in rodents has been shown to effectively model several of the behavioral and endocrine impacts of ACEs and has been utilized to investigate the underlying mechanisms contributing to disease. However, the precise neural mechanisms responsible for mediating the impact of ELS on vulnerability to psychiatric illnesses remain largely unknown.

**Methods:** We use behavior, immunoassay, *in vivo* LFP recording, histology, and patch clamp to describe the effects of ELS on stress behaviors, endocrinology, network states, protein expression, and cellular physiology in male and female mice.

**Results:** We demonstrate that a murine maternal separation (MS) ELS model causes sex-dependent alterations in behavioral and hormonal responses following an acute stressor. Local field potential (LFP) recordings in the basolateral amygdala (BLA) and frontal cortex (FC) reveal similar sex-dependent alterations at baseline, in response to acute ethological stress, and during fear memory extinction, supporting a large body of literature demonstrating that these network states contribute to stress reactivity and vulnerability to psychiatric illnesses. Sex differences were accompanied by altered physiology of BLA principal neurons in males and BLA PV interneurons in females.

**Conclusions:** Collectively, these results implicate novel, sex-dependent mechanisms through which ACEs may impact psychiatric health, involving altered cellular physiology and network states involved in emotional processing.

## Introduction

Adverse childhood experiences (ACEs) dramatically increase vulnerability to psychiatric illnesses (VanTieghem & Tottenham, 2018), although the complete view of the neuronal mechanisms underlying this relationship remain unclear. Psychiatric illnesses have been associated with changes in the activity within and across the frontal cortex(FC)-amygdala circuit (Chai et al., 2011; Liu et al., 2015; Reincke & Hanganu-Opatz, 2017; Tye et al., 2011); however, the impact of early life stress (ELS) on FC-amygdala network states has not been well characterized. Recent work from our laboratory has demonstrated that chronic stress paradigms in adulthood induced long-term alterations in network states in the basolateral amygdala (BLA) associated with behavioral deficit (Walton, Najah et al., 2023). Here we investigate the impact of early life stress on FC-BLA network and behavioral states. Improving our understanding of how the brain switches between adaptive and maladaptive states may reveal novel targets for treating these psychiatric illnesses.

Glucocorticoids and catecholamines released at the hypothalamic-pituitary-adrenal (HPA) axis terminal are the primary effectors through which stress is signaled internally, often resulting in behavioral changes. Despite this ubiquity of the HPA axis in the stress response, stress is highly heterogeneous in terms of its triggers, subjective effects, and neural consequences. For example, acute, chronic, and social stress appear to differentially impact the brain and behavior (Cabib et al., 1988; McEwen, 2017; McEwen et al., 2015; Motta & Canteras, 2015), with effects appearing particularly salient when the stressor occurs in development (Bingham et al., 2011; Eiland & Romeo, 2013; S. Xie et al., 2021). Moreover, behavioral strategies for responding to the same stressor can be classified as either adaptive or maladaptive depending on the context in which they occur and the degree to which they impact allostatic load (Bourdon et al., 2020; Ferguson et al., 2000). Highly impactful stressors – such as neglect and resource scarcity – increase engagement with risky behaviors as well as vulnerability to psychiatric disorders in some individuals, while others appear resilient to these effects (Campbell et al., 2016; Felitti et al., 1998). As such, effective animal models of ELS should be leveraged to investigate the mechanisms contributing to these disparate phenotypes.

While these models have faced criticism for producing inconsistent results due to varying design and stress intensity, these discrepancies may be reflective of the heterogeneity of ethological stressors and their distinct impacts on internal states across underexplored variables. For example, maternal separation (MS) models have been shown to drive anxiety-like behaviors in adult males while limited bedding and nesting (LBN) models drive this phenotype in adult females (Demaestri et al., 2020). Similarly, MS has been shown to increase the expression of genes associated with corticotropin-releasing hormone (CRH) ligand and receptor synthesis in the amygdala of males, while LBN shows similar effects in females (Demaestri et al., 2022). These results suggest that developmental stressors relating to resource scarcity may be more salient in females, while males are more impacted by those resulting from atypical parental care. In accordance with this emerging narrative, we demonstrate that MS impacts stress hormone levels and behavior in a subset of males, with no effects on these parameters in females. Furthermore, we investigate the impact of MS on BLA and FC network oscillations under basal conditions and in association with distinct conditioned and ethological stressors, addressing issues concerning stress heterogeneity.

Functional magnetic resonance imaging (fMRI) studies in humans have demonstrated that developmental stress impacts functional connectivity between the FC and amygdala, as well as reactivity to threat in both regions (Gee et al., 2013; Javanbakht et al., 2015). While similar effects have been postulated in animal models, there is a surprising lack of translational research defining the specific effects of ELS on this circuit. LFPs influence routing of information through the brain and represent a high-resolution metric for observing local network states (Akam & Kullmann, 2010; Davis et al., 2017; Salinas & Sejnowski, 2001). PV interneurons in the BLA are developmentally regulated, appear to be impacted by developmental stress (Abraham et al., 2023; Manzano Nieves et al., 2020), and have been shown to influence network states corresponding with changes in behavior (Antonoudiou et al., 2022; Davis et al., 2017; Ozawa et al., 2020).Therefore, we hypothesized that MS would impact BLA PV expression and electrophysiology contributing to altered behavioral states and FC-BLA network states.

Our results confirm our hypothesis that ELS impacts theta oscillations in the FC-BLA circuit at rest and in response to ethological and conditioned stressors. These findings are consistent with human literature implicating this circuit. Surprisingly, we found that MS did not impact the expression of PV or an associated class of developmentally regulated plasticity-limiting lectins called perineuronal nets (PNNs) in the BLA. Whole cell recordings revealed that ELS did not impact BLA PV electrophysiology in adult males, however principal neurons displayed reductions in peak firing rate as well as shifted resonance properties. Conversely, ELS in females did not impact principal neuron physiology, but resulted in PV interneurons with shifted resonance properties. Collectively, these effects suggest novel mechanisms through which ELS appears to alter network states, with important implications for sex differences in the expression of psychiatric illnesses. While further research on this topic is required, identifying the impact of ELS on network states in the FC-BLA circuit could provide a useful biomarker and target for treating psychiatric vulnerability following ACEs.

## Methods

### Subjects

All mice were raised in the Tufts University animal research facilities. Dams were ordered timed-pregnant from Jackson Laboratories or bred in-house. All procedures were conducted in accordance with Tufts University Institutional Animal Care and Use Committee (IACUC) guidelines. C57BL/6J mice were used for all experiments in which genetic identification of PV was not necessary, excluding a subset of behavioral experiments performed on DBA/2J mice to test for genetic contributions guiding diverse behavioral responses. Genetic identification of PV for electrophysiology was achieved using a transgenic C57BL6 line which expresses tdTomato under the *Pvalb* promoter (C57BL/6-Tg(Pvalb-tdTomato)15Gfng/J) or a cross between 2 strains that are congenic on the C57BL6 genetic background and also express tdTomato in PV interneurons (B6.129P2-*Pvalb^tm1(cre)Arbr^*/J & B6.Cg-*Gt(ROSA)26Sor^tm9(CAG=tdTomato)Hze^*/J). All experiments used both male and female mice that were raised in our colony and were exposed to 3 hrs of maternal separation for 15 days during the 1^st^ 3 weeks postnatal and or reared in standard housing conditions as controls.

### LFP surgeries

Custom headmounts for LFP recordings were fabricated by soldering PFA-coated silver wire (coating removed at contact points) to an EEG/EMG headmount (Pinnacle Technology, cat. # 8201). Mice were intraperitoneally injected with 100 mg/kg ketamine mixed with 10 mg/kg xylazine for anesthetization and subcutaneously injected with 0.5 mg/kg Buprenorphine Extended-Release Lab formulation (ZooPharm) for analgesia. Mice were placed in a murine-specific stereotaxic apparatus, the scalp was shaved, disinfected using iodine and 70% ethanol, and a lengthwise incision was made to expose the skull. The skull was leveled along the anterior-posterior and medial-lateral axes and a burr hole was created over the right hemisphere through which the silver wire electrode of the prefabricated headmount was implanted into the BLA (AP −1.5 mm, ML +3.3 mm, DV −4.5 mm). The headmount was fixed to the skull using three 0.12” EEG mouse screws (Pinnacle Technology, cat. # 8212) which served as a lead in the PFC, a reference electrode, and a ground electrode. Bone cement was used as surgical dressing and to provide structural support. Mice were each carefully monitored for 3 days and network recordings were not initiated for ≥ 3 weeks post-surgery.

### Behavior

Juvenile (4 weeks postnatal) and adult (10 weeks postnatal) mice were observed for behavioral changes independent of LFP recording. Behavioral paradigms involving LFP recordings were performed between 15 – 20 weeks postnatal and are discussed in the ‘network recordings’ section below. For all behavioral paradigms, mice were acclimated to the experimental room for ≥ 1 hr, the experimental chamber was thoroughly disinfected with 70% ethanol between trials, and order was counterbalanced. Mouse location and movement were detected using ethovision (Noldus) or custom-written python scripts when specified.

#### Open Field Test

Each mouse was placed in the center of a square 406 x 406 x 380 mm chamber. Once detected, a 10 min recording was initiated. Automated behavioral detection measured the percentage of time spent in the center (230 x 230 mm) versus the periphery.

#### Elevated Plus Maze

Each mouse was placed in the center of an elevated plus maze (EPM). The dimensions of the center of the EPM were 62 x 62 mm, while each arm measured 350 x 62 mm with a closed-arm wall height of 152 mm. Once detected by the software, a 10 min recording was automatically initiated. Automated behavioral detection measured the percentage of total time spent in the open arms.

#### Forced swim test

Each mouse was placed into the center of a cylindrical forced swim test (FST) chamber with a 216 mm diameter and a 250 mm height filled with room temperature water for a 6 min test. Movement was detected using a custom script Python script which uses machine learning to detect limbs and measure mobility, and which produces results that are consistent with human-scored data. Automated behavioral detection quantified both the percentage of total time spent immobile as well as the latency to display immobility from the beginning of the trial.

#### Social Interaction Test

Each mouse was placed in the center chamber of a 3-chamber social interaction apparatus. Each square chamber measured 280 x 280 x 350 mm. Each outer chamber contained a small wire cage (105 mm height x 100 mm base diameter) in its center, under which a resident mouse was placed, such that they were able to interact with the experimental mouse and exchange vocal and olfactory signals. To familiarize each experimental mouse with the apparatus – and to test for innate preference – an initial 5 min trial was recorded, during which they were allowed to explore the arena in the absence of a resident mouse. Subsequently, after thoroughly disinfecting the apparatus with 70% ethanol, a novel age- and sex-matched resident mouse was placed under the cage in one of the two outer chambers. The experimental mouse was then placed back in the center of the arena, and an additional 10 min experimental recording was collected to test for the impact of ELS on social interaction in males and females. The outer chamber in which the resident mouse was placed was counterbalanced. Automated behavioral detection quantified the percentage of total time spent in the chamber containing the novel resident.

### Network recordings

LFPs were recorded at baseline and in response to acute and conditioned stressors from single-housed, tethered C57BL/6J mice between 15-20 weeks postnatal. Each mouse was given ≥ 1hr to acclimate to the room before testing. During data collection, mice were allowed to move freely throughout the cage. Data were acquired through LabChart (ADInstruments) at 1 or 4 KHz and amplified x100. Data were processed in Python using SAKEverse, an open-source platform for spectral analysis of LabChart recordings. Signal quality was verified by visually inspecting power spectral density (PSD) features, and low-quality recordings were removed. Mains noise was removed between 59-61 Hz and replaced using the piecewise cubic hermite interpolating polynomial method (PCHIP). Datapoints with an absolute value exceeding 7x the median were removed as outliers (60 s window) and replaced with the nearest bin. For network recordings, each brain region was analyzed independently. Biologically relevant frequency bands were defined as 2 – 5 Hz (low theta), 6 – 12 Hz (high theta). Power within these bands was calculated the mean across all included frequencies.

#### Baseline

Baseline recordings were collected across a 24 hr period. Subjects were tethered with an LFP preamp and placed in a pinnacle cage (203 mm x 254 mm diameter) equipped with standard bedding and provided with *ad libitum* access to food/water. Each mouse was allowed to acclimate to this environment overnight before any recording was initiated. Fast Fourier transform (FFT) was performed on the recording with a window of 5 s and an overlap of 0.5 s. Frequencies between 2 - 150 Hz were analyzed, and the power at each frequency was normalized to the mean power within this range before binning across frequency bands, to control for possible differences in preamplifier processing. Data were then averaged across time. Low- to high-theta ratio was defined as the mean power across low theta divided by the mean power across high theta for each animal.

#### Fear conditioning

Contextual fear conditioning experiments were designed based off previous reports from our laboratory (Davis et al., 2017). The conditioned context was a cubic chamber with a metal grid floor measuring 255 x 285 x 290 mm (measurements without grid floor: 255 x 285 x 325 mm). Briefly, on day 1, mice were subjected to a baseline trial lasting 500 s followed by three conditioning trials separated by 3 hrs. Each conditioning trial lasted 500 s and consisted of 2 s, 0.70 mA foot shocks at 198 s, 278 s, 358 s, and 438 s. On days 2 and 3, mice were subjected to 4 extinction trials per day during which no shocks were administered, each lasting 1200 s and separated by 2 hrs. On day 4, mice were subjected to a final 500 s no-shock retrieval trial. Mice were tethered during each trial, however recordings were not collected during conditioning trials due to electrical noise. Videos were acquired using FreezeFrame (Actimetrics), but, due to artifacts from the tether, were exported and movement was detected using a custom python script which tracks the darkest spot on the video. To account for changes in immobility resulting from trials of varying length, behavioral quantification was limited to the first 450 s of each trial. Pixel-based thresholding of freezing was manually calibrated and uniformly applied across subjects to obtain behavioral results that were largely consistent with previous reports using this paradigm (Davis et al., 2017). Subjects were excluded from analyses if the corresponding video file from any of the analyzed timepoints had been corrupted (i.e. all subjects were represented at all timepoints).

FFT was performed on the recording with a window of 1 s and an overlap of 0.5 s. Each post-conditioning recording was normalized, within-subject and within-band, to the average power of the baseline recording. For preliminary analyses, we isolated the network state during freezing vs. exploratory behavior, however we did not observe substantial differences and ultimately we elected to analyze these data collectively.

#### Predator odor

Tethered mice habituated to the environment for 24 hrs before recording was initiated. After 1 hr of baseline recording, a lab tissue containing 16 drops (potent but not saturated) of bobcat urine (predator odor) was introduced to the cage and an additional 1 hr recording was collected. FFT was performed on the recording with a window of 5 s and an overlap of 0.5 s. Power from the baseline period was averaged across time, and power during treatment was represented as % of average baseline power at that frequency band, which was subsequently averaged across 10 min bins.

#### Looming disk

The looming disk chamber was composed of a rat cage (375 x 285 x 220 mm) with a 3D-printed triangular shelter (100 x 120 x 120 mm) in one corner. A computer monitor (300 x 535 mm) connected to a raspberry pi and displaying a solid white screen was placed over the cage. The monitor screen was larger than the rat cage, however care was taken to ensure that the looming disk stimulus was fully visible and centered over the arena. 1-2 days prior to testing, each mouse experienced a 20 min habituation trial during which it was tethered and allowed to freely explore, however no stimuli were presented. On the test day, each mouse was again tethered and placed in the arena. No stimuli were presented during the first 3-5 min, after which a black circle was manually triggered to appear and rapidly expand at a time when the mouse was outside of the shelter. Care was taken to trigger the disk when the mouse was mobile and the head was tilted upwards to ensure that the stimulus was detected. Each mouse experienced 5 stimulus repetitions separated by > 1 min. Behaviors were manually scored from video. Generally, tail rattle and darting were preceded by a brief (< ∼1 sec) bout of freezing – in such cases, the active behavior was scored. Lighting issues or camera placement rendered the behavior unquantifiable in a subset of videos – in this case, or in the case that the mouse showed no discernable behavioral changes following stimulus onset, the corresponding stimulus presentation was excluded from analyses.

FFT was performed on the recording with a window of 1 s and an overlap of 0.5 s. Baseline power for each frequency was calculated as the average power across minutes 2-3 of habituation (day 2). For measuring changes in LFP during disk presentation, power was averaged across the 10 s stimulus and normalized to baseline. Data were then averaged across biologically relevant frequencies and compared to behavioral coping strategies.

### Blood collection & corticosterone measurement

Corticosterone (CORT) concentration was measured from serum. *Sample collection.* Blood was collected between ZT 14-15 using either submandibular bleed (baseline) or rapid decapitation (post-restraint). Baseline and post-restraint measures were both collected for all subjects. In the case of post-restraint measurements, mice were individually placed in custom restraint devices for 30 min (fabricated by introducing an air hole to a 50 mL Falcon tube) and subsequently individually housed for an additional 30 min – which is within the temporal range of the peak CORT response (Dopfel et al., 2019) – before blood sample collection. Samples remained on ice during collection. To extract serum, samples were taken off ice (room temperature) for 20 min and subsequently centrifuged at 1.8 x g for 15 min at 4°C. *Corticosterone measurement*. CORT concentration was quantified using a CORT enzyme-linked immunosorbent assay (ELISA) kit (Enzo Life Sciences, cat. # ADI-900-097). Serum samples were run in duplicates of 5 uL, and results were interpolated against a standard curve of known concentration. The normalized CORT response was calculated with the formula (restraint – baseline)/(restraint + baseline).

### Immunohistochemistry

Mice were sacrificed using rapid decapitation with isoflurane anesthetization and brains were drop-fixed overnight in 4% paraformaldehyde at 4°C. Subsequently, brains were serially cryoprotected in 10% and 30% sucrose (percentage by weight) in phosphate buffered saline (PBS), snap-frozen, and stored at - 80°C. Coronal sections were collected at 25 μm. For staining, sections were first washed 3 times (10 min, room temperature) in PBS, then incubated in a blocking solution containing 10% normal goat serum in PBS with 0.3% Triton X-100 (1 hr, room temperature). Next, sections were incubated in a solution containing a monoclonal primary antibody raised against parvalbumin (Sigma-Aldrich cat. # P3088) at a concentration of 1:1000 in blocking solution (72 hr, 4°C). Following 3 additional washes in PBS (10 min, room temperature), sections were incubated in a solution containing both anti-Rabbit Alexa Fluor 647 (Invitrogen, cat. # A-21245) at a concentration of 1:200 and a fluorescent-conjugated lectin (WFA) which binds carbohydrate structures and excites at 495 nm (Vector Laboratories, cat. # 1351) at a concentration of 2 ng/mL in blocking solution (2 hr, room temperature) under light-deprived conditions. Finally, sections were washed in PBS 3 more times (10 min, room temperature) and mounted to microscope slides using VECTASHIELD Antifade Mounting Medium with DAPI (Vector Laboratories, cat. # H-1200-10). Sections were imaged on a Nikon A1R confocal microscope at the Tufts University Center for Neuroscience Research Microscopy Core. Z-stacks were collected using slice thickness of 1-2 µm, with upper and lower bounds set such that the full 25 µm depth was captured. All images were acquired using a 20X objective. Analyses were performed in ImageJ. For quantification, a maximum intensity projection was applied to each image. Each mouse was treated as an independent datapoint, and cell counts for each were averaged across 5-6 distinct, matched unilateral sections containing BLA. Therefore, reported values should not be influenced by hemispheric or BLA subregion-specific changes in expression.

### *Whole cell* electrophysiology

Mice were sacrificed using rapid decapitation under isoflurane anesthesia. Brains were rapidly extracted and submerged in ice cold cutting solution containing (in mM): 150 sucrose, 15 glucose, 33 NaCl, 25 NaHCO_3_, 2.5 KCl, 1.25 NaH_2_PO_4_, 1 CaCl_2_, 7 MgCl_2_ (300 – 310 mOsm). 350 μm coronal sections were prepared using a vibrating microtome and incubated at 33°C in standard artificial cerebrospinal fluid (aCSF) containing (in mM): 126 NaCl, 10 glucose, 2 MgCl_2_, 2 CaCl_2_, 2.5 KCl, 1.25 NaHCO_3_, 1.5 Na-pyruvate, 1 L-glutamine (300 – 310 mOsm) for ≤ 1 hr before experimentation. During recording, slices were continually perfused at a rate of 1.799 mL/min with 33°C aCSF – maintained using an in-line heater (Warner Instruments) – saturated with 95% O_2_/5% CO_2_. Current clamp recordings were collected from visually identified principal neurons and fluorescently labelled PV interneurons (PV-tdTomato) using a 200B Axopatch amplifier (Molecular Devices), LabChart 8 acquisition software (AD Instruments), and micropipettes fabricated from borosilicate glass (World Precision Instruments) with a tip resistance of 5-7 MΩ which were backfilled using an internal solution containing (in mM): 130 K-gluconate, 10 KCl, 4 NaCl, 10 HEPES, 0.1 EGTA, 2 Mg-ATP, 0.3 Na-GTP (7.25 pH, 280 – 290 mOsm/L H_2_O). Input-output curves and rheobase were generated from 0 – 150 pA current injections in 10 pA steps. Resonant activation frequencies were determined using a 3s suprathreshold sinusoidal chirp ranging from 6 – 60 Hz. Representative waveforms were averaged across the first 3 action potentials elicited at each step from rheobase to rheobase + 20 pA. Intrinsic electrophysiological characteristics such as input resistance, impedance, and whole cell time constant were assessed for each neuron. Cells were excluded from the analysis if their series resistance exceeded 20 MΩ or varied by more than 20% during the experiment.

### Analyses

#### Statistical analyses

Statistics were computed in Prism 10. Outliers were removed from each dataset using the ROUT test with Q = 1%. For column analyses, unpaired student’s t-tests were used barring cases where the F-test was significant (< 0.05), in which case statistics were re-run using Welch’s correction. To test for genetic contributions to opposing behavioral phenotypes in males in the SI test, data were analyzed using a 2-way ANOVA with Fisher’s LSD. Both timeseries and freezing datasets were analyzed using mixed effects modeling (to handle missing data resulting from outlier removal) with Šidák multiple comparisons corrections to compare stress condition with time bin or treatment. In all cases, data were only considered significant with a P value of < 0.05.

## Results

Control (CTL) and ELS male mice were observed during low and high stress behavioral tests at both late adolescence and early adulthood (4 & 10 weeks postnatal). In the FST, ELS mice demonstrated an increased latency to immobility as well as reduced percentage time immobile at both time points (Fig. 2A-D). This corresponded with a reduced CORT response following acute stress in adults (Fig. 2E). Interestingly, while ELS did not significantly influence social behavior at either time point when measured using a Student’s t test, the corresponding p-value was 0.0500 and a subsequent ANOVA testing for genetic contributions to the impacts of ELS on SI between male C57BL/6J and DBA/2J mice revealed a significant effect of stress (DBA table). Collectively, these results indicate the effects of ELS on SI merit further investigation. ELS mice were not different from CTL in the low stress open field test (Fig. 2 H-I) or EPM (Fig. 2 J-K). Female mice did not demonstrate any significant behavioral or endocrine effects of ELS (Supp. Fig. 1). To look for genetic mediators of the adverse impacts of ELS we examined behavior in a separate genetic strain of mice (DBA/2J). Like C57BL/6J mice, DBA2/J mice demonstrated no significant differences in stress behaviors in the open field test or the social interaction test resulting from ELS (Supplemental Figure 2). When male CTL and ELS mice were compared across genetic background, however, a significant effect of strain was observed in ELS mice (DBA table).

**Figure 1.**
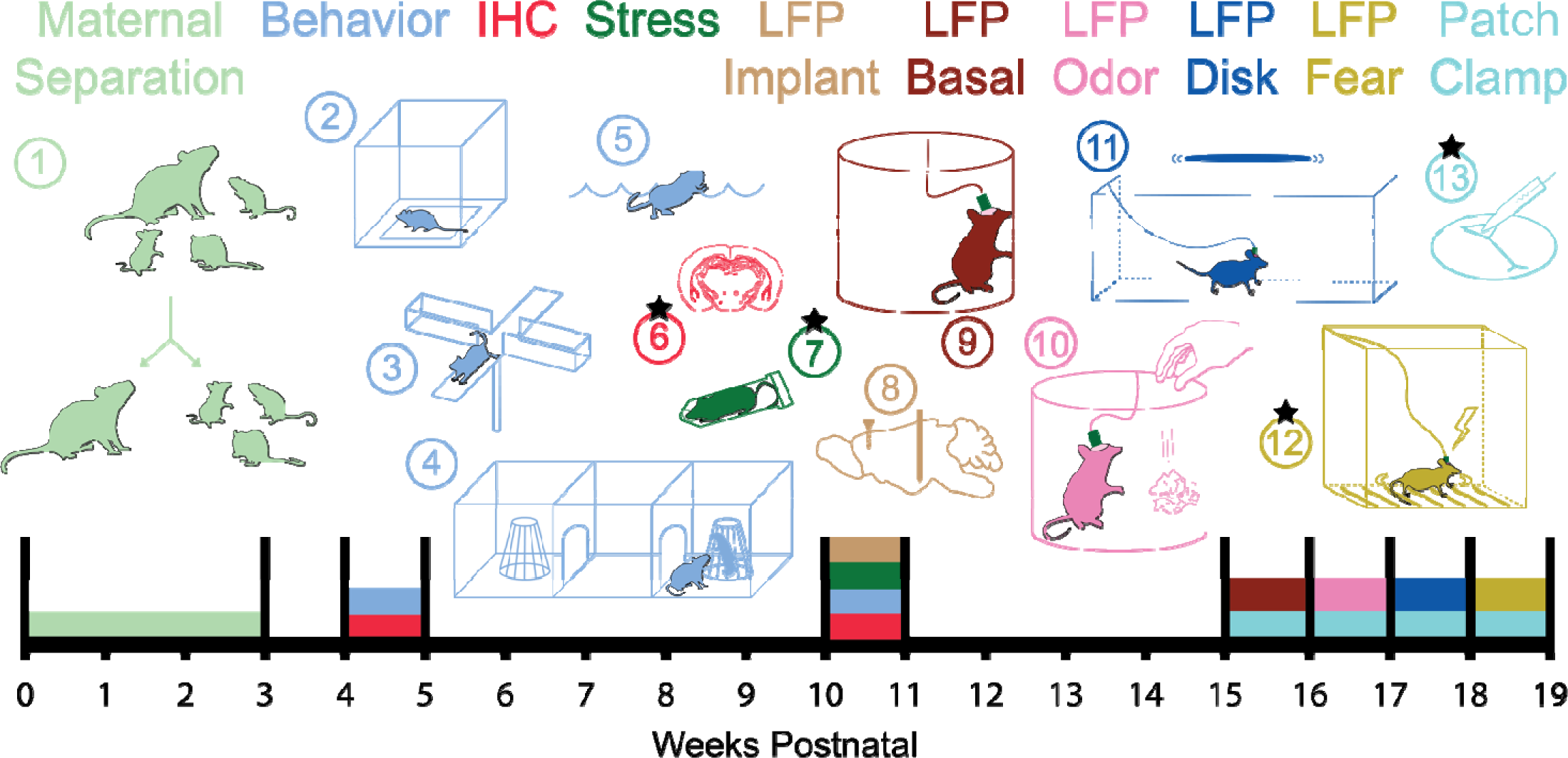
Timeline of MS intervention and experimentation.

**Figure 2.**
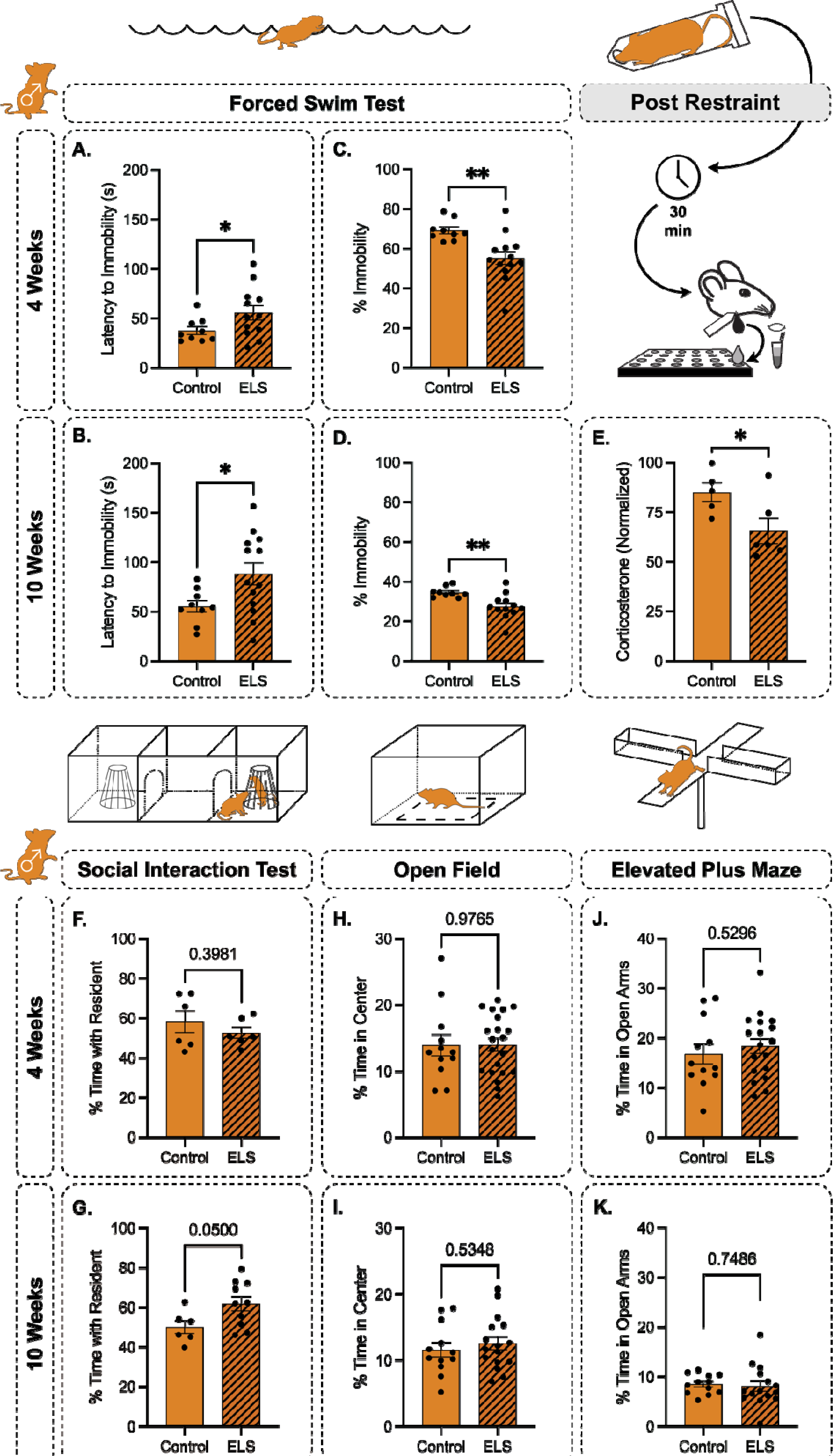
ELS increases escape behavior in the FST and reduces post-stress CORT in males with no impact on social or exploratory behaviors. Male ELS mice display significantly increased latency to immobility in the FST at 4 weeks (**A**) and 10 weeks (**B**) postnatal, as well as reduced percentage of total time spent immobile at 4 weeks (**C**) and 10 weeks (**D**). At 10 weeks postnatal, male mice display a reduction in the CORT response following acute restraint stress (**E**). Male mice showed no differences in the percentage of total time spent with a resident in the SI test at 4 weeks (**F**) or 10 weeks (**G**), in the percentage of total time spent in the center vs. the periphery in the open field test at 4 weeks (**H**) or 10 weeks (**I**), or in the percentage of total time spent in the open arms in the elevated plus maze at 4 weeks (**J**) or 10 weeks (**K**).

**Supplemental Figure 1.**
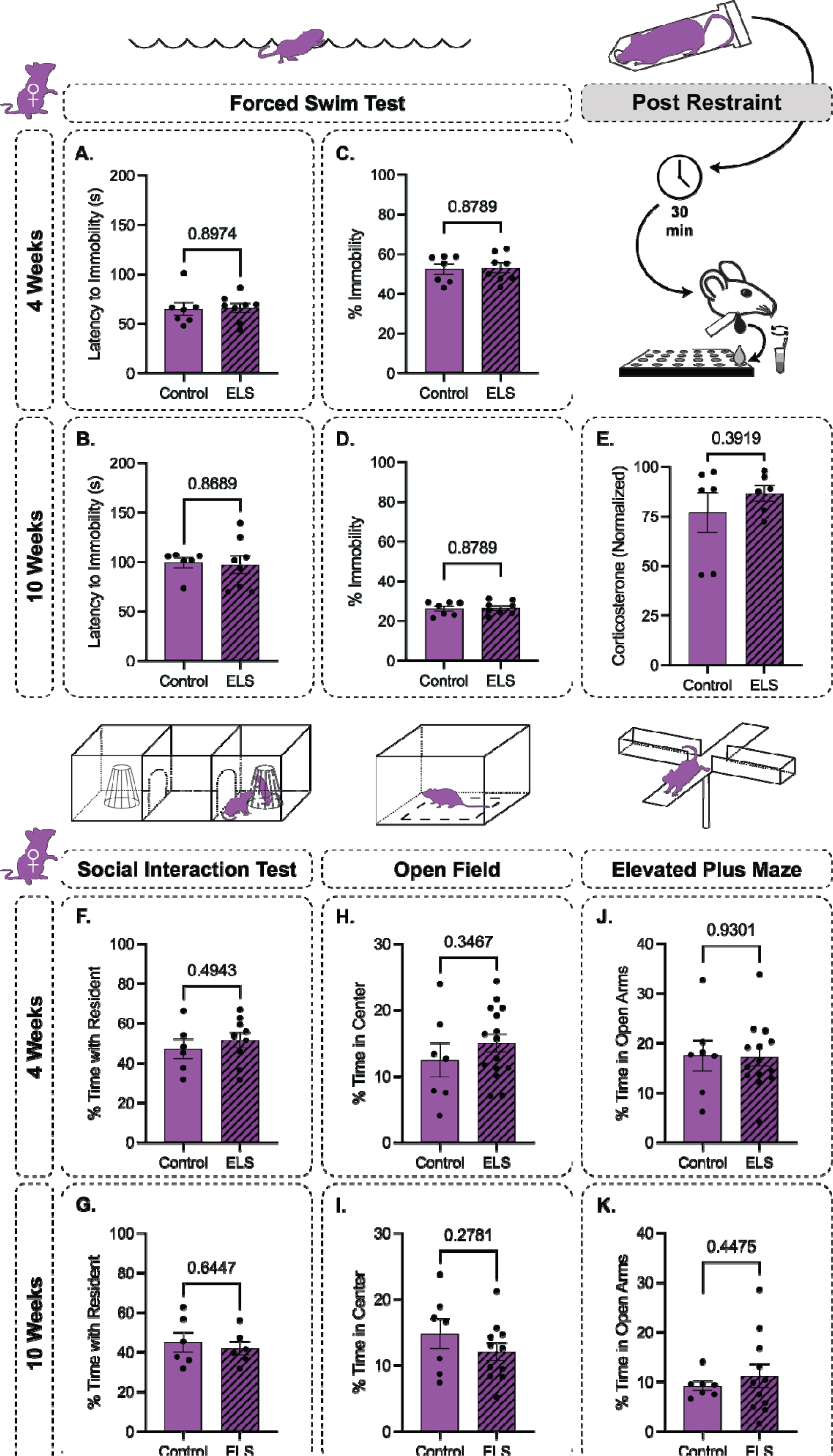
ELS does not impact escape behavior in the FST, post-stress CORT, social behavior, or exploratory behaviors in females. Female ELS mice do not display any differences in latency to mobility during the FST at 4 weeks (**A**) or 10 weeks (**B**) postnatal, and are also not significantly different in terms of percentage of total time spent immobile at 4 weeks (**C**) or 10 weeks (**D**) postnatal. Female ELS mice are not significantly different in the CORT response following an acute restraint stress (**E**), in the percentage of total time spent with a resident in the SI test at 4 weeks (**F**) or 10 weeks (**G**) postnatal, in the percentage of total time spent in the center vs. the periphery during the open field test at 4 weeks (**H**) or 10 weeks (**I**) postnatal, or in the percentage of total time spent in the open arms of the elevated plus maze at 4 weeks (**J**) or 10 weeks (**K**) postnatal.

**Supplemental Figure 2.**
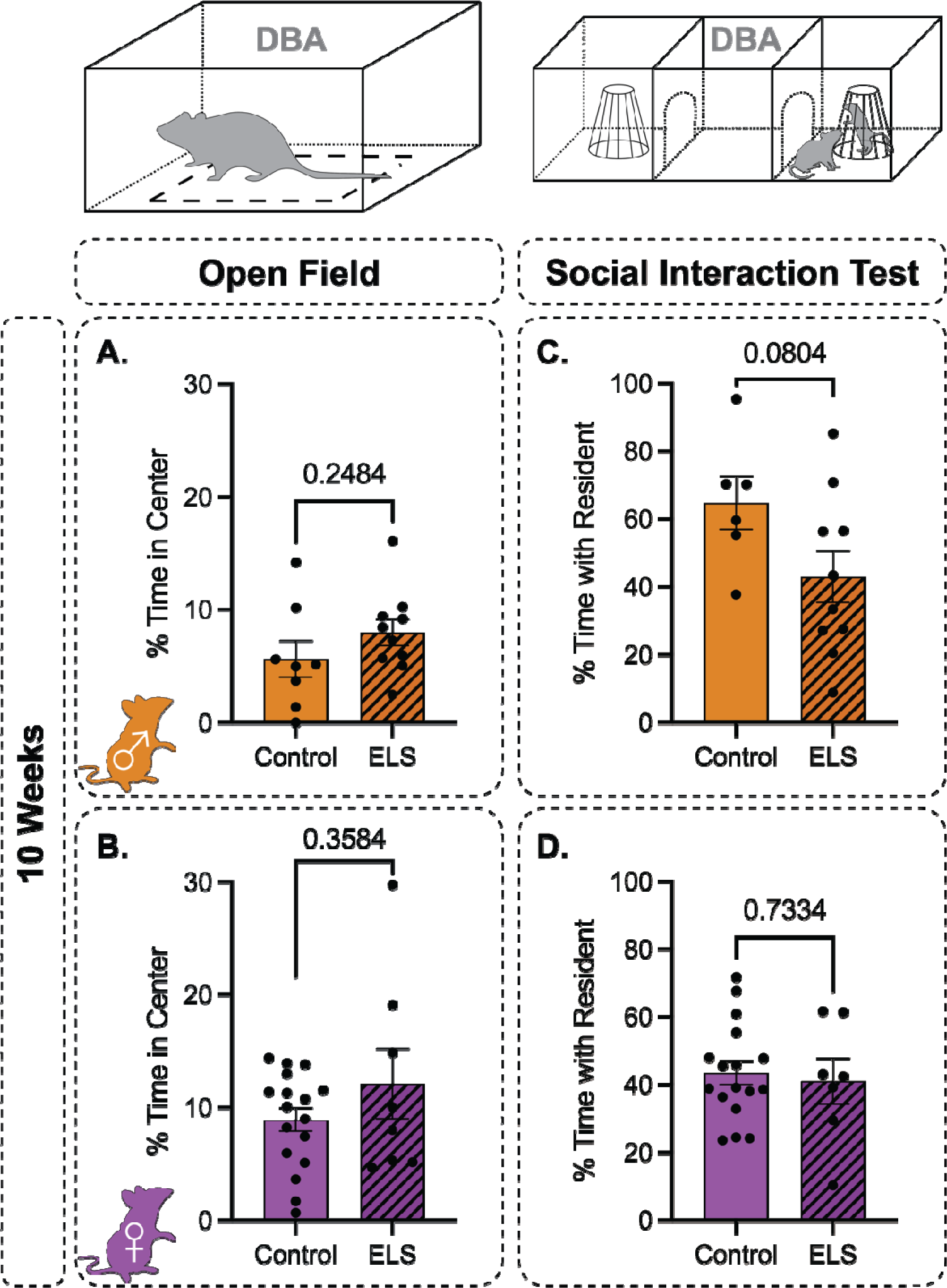
ELS does not impact exploratory or social behavior in males or females. DBA/2J ELS mice are not different from CTL in percentage of total time spent in the center vs. the periphery of the open field test in males (**A**) or females (**B**). Similarly, no significant differences are observed in terms of the percentage of total time spent with a resident in the SI test in males (**C**) or females (**D**).

Next, we looked for changes in network states following ELS by measuring LFP power at rest across frequency ranges which have previously been associated with processing stressful stimuli. Males did not demonstrate any significant effects of ELS on baseline LFP power in the ranges of low theta or high theta, or in the low-to-high theta ratio, in the BLA (Fig. 3B-D) or FC (Fig. 3F-H). Females also showed no significant changes in the baseline LFP resulting from ELS across these categories in the BLA (Supp. Fig. 3B-D) or in low theta in the FC (Supp. Fig. 3F), however a significant reduction in high theta power (Supp. Fig. 3G) and a significant increase in the low-to-high theta ratio (Supp. Fig. 3H) were observed in the FC.

**Figure 3.**
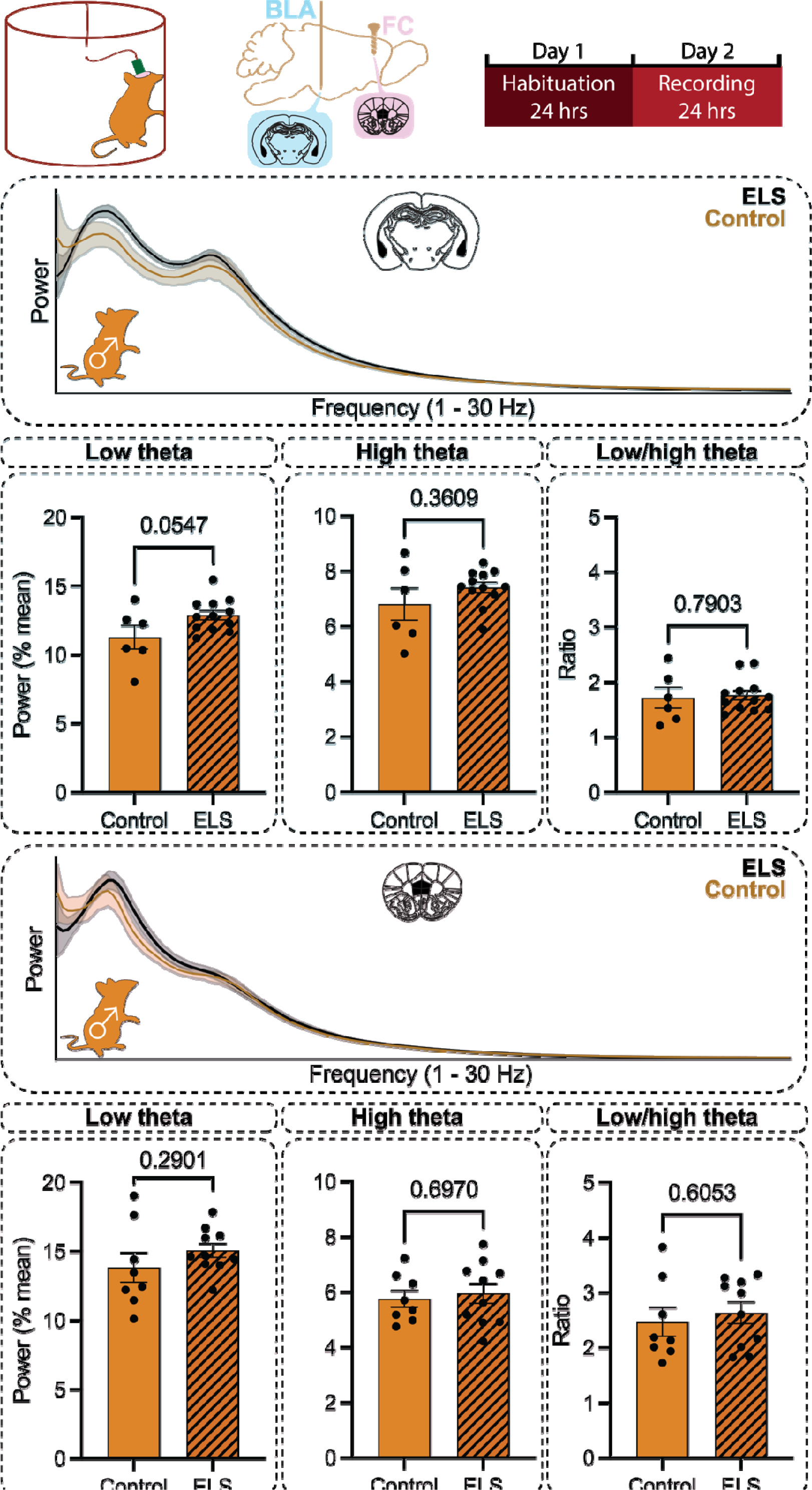
ELS does not impact theta oscillations in the BLA or FC in males at rest. Male ELS mice display LFPs in the BLA which are not significantly different from CTL at rest in the range of low theta (**B**), high theta (**C**), or the low-to-high theta ratio (**D**). Similarly, LFPs in the FC are not significantly different from CTL at rest in the range of low theta (**F**), high theta (**G**), or the low-to-high theta ratio (**H**).

**Supplemental Figure 3.**
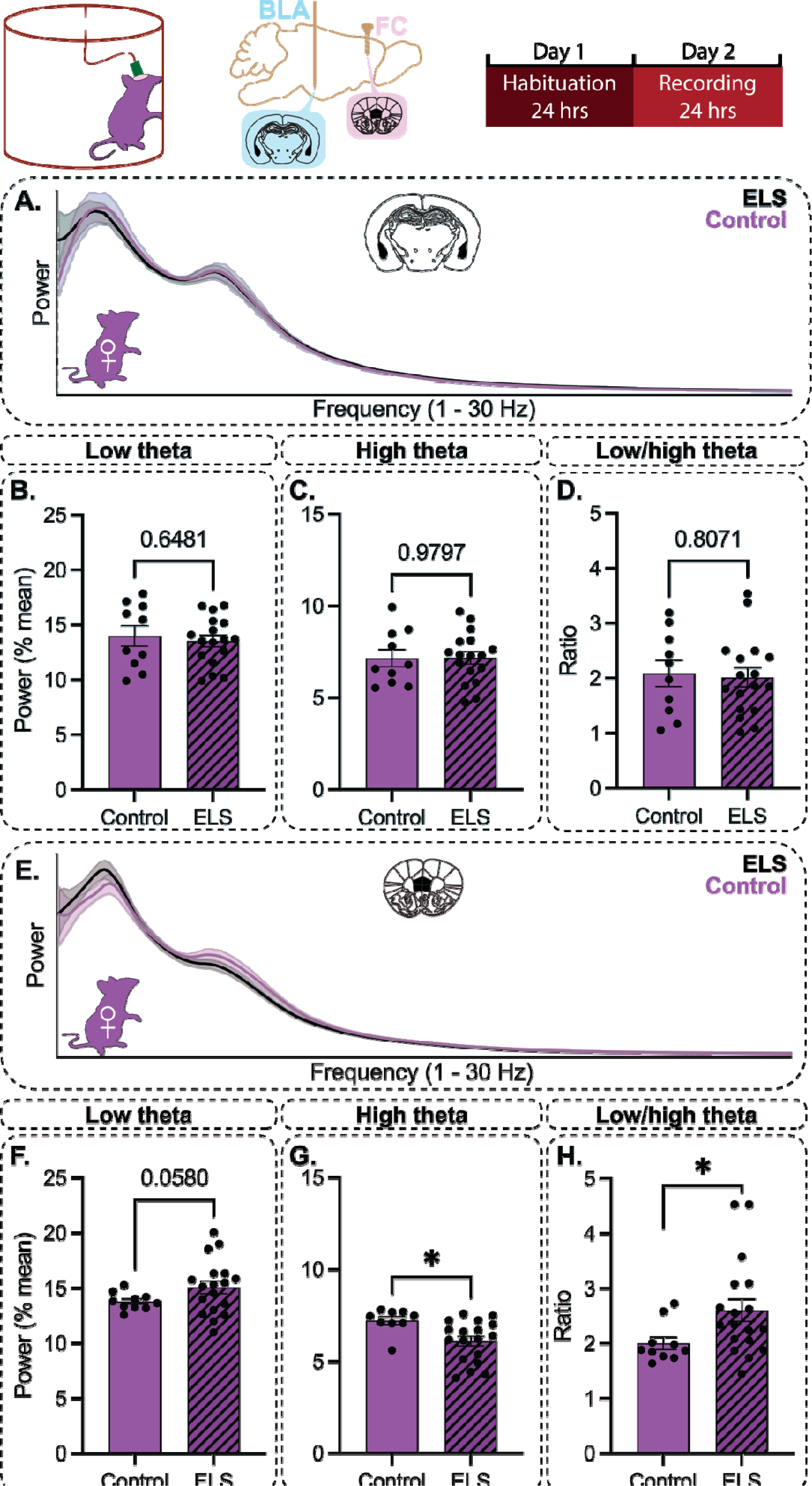
ELS increases the ratio of low-to-high theta in the FC of females at rest. Female ELS mice display LFPs in the BLA which are not significantly different from CTL at rest in the range of low theta (**B**), high theta (**C**), or the low-to-high theta ratio (**D**). In the FC, while LFPs are not significantly different from in the range of low theta (**F**), ELS females display a significant reduction in high theta power (**G**) and a significant increase in the ratio of low-to-high theta (**H**).

The lack of observed changes in the baseline LFP following ELS was expected due to the dynamic nature of these oscillatory states associated with specific behavioral states. Thus, to investigate the impact of ELS on network state changes contributing to altered behaviors resulting from ELS, we subjected CTL and ELS mice to fear conditioning and extinction while observing LFP power. Male mice demonstrated a significant effect of post-conditioning trial on freezing behavior with no difference between CTL and ELS groups (Fig. 4A). Consistent with this observation, male ELS mice did not display any significant LFP differences in BLA low theta or high theta shifts from pre-conditioning baseline during the first trial on the first day of extinction (Fig. 4B-C), during the first trial on the second day of extinction (Fig. 4D-E), or during a final recall trial (Fig. 4F-G). There were also no significant changes observed across these trials and frequencies in FC LFP power (Fig. 4H-M). Female mice also demonstrated a significant effect of post-conditioning trial on freezing behavior. Female ELS mice were not significantly different from CTL in freezing behavior (Supp. Fig. 4A) or in BLA low or high theta shifts from baseline during the first trial on the first day of extinction (Supp. Fig. 4B-C) or the first trial on the second day of extinction (Supp. Fig. 4D-E). During recall, female CTL and ELS mice were not significantly different in terms of their low theta changes in power (Supp. Fig. 4F), however high theta power was significantly reduced from baseline in CTL mice relative to ELS (Supp. Fig. 4G). There were no significant differences observed in the FC (Supp. Fig. 4H-M).

**Figure 4.**
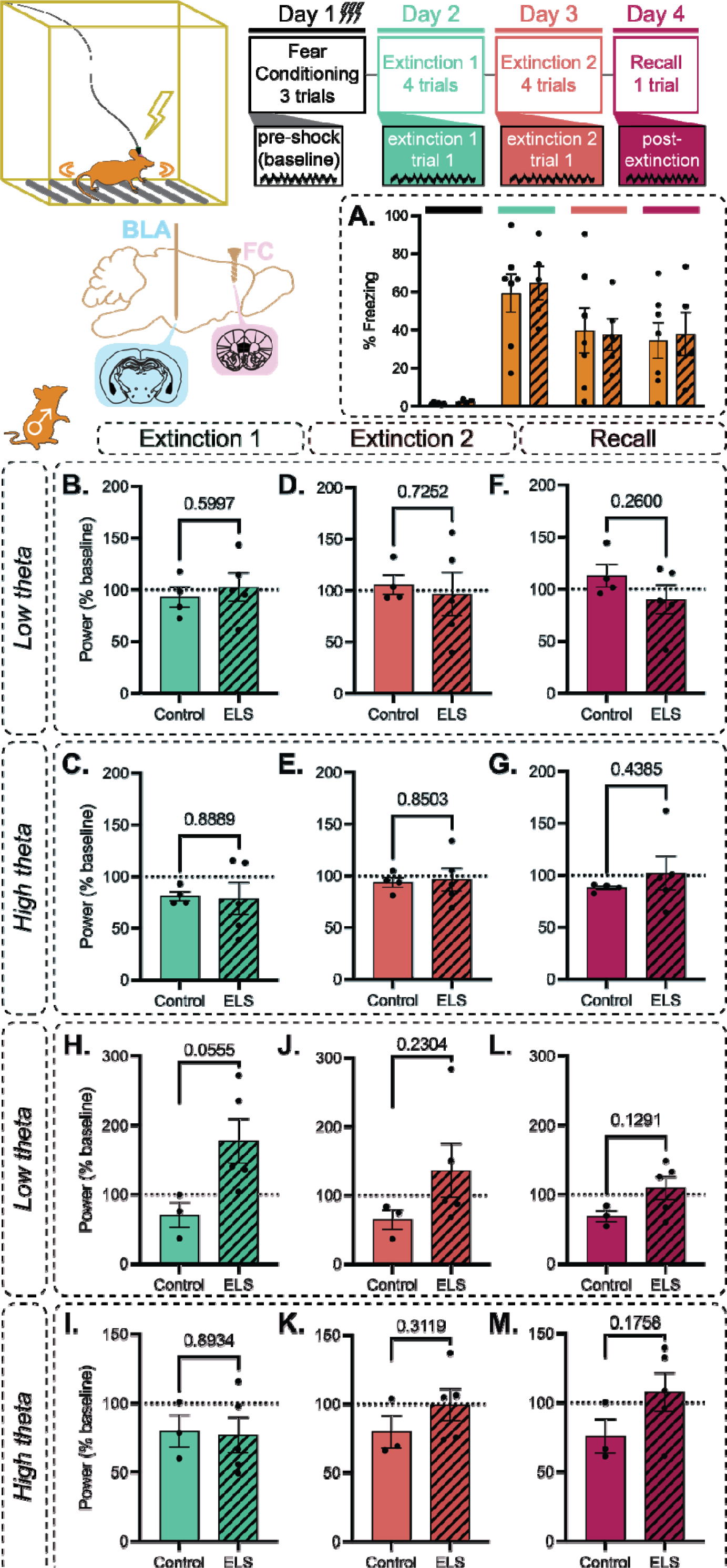
ELS does not influence freezing behavior or network states in the BLA and FC during fear conditioning and extinction in males. Male ELS mice are not significantly different from CTL in terms of freezing behavior across fear conditioning and extinction trials (**A**). In the BLA, during the 1^st^ extinction trial on the 1^st^ extinction day there are no differences observed in low theta (**B**) or high theta (**C**) network responses, during the 1^st^ extinction trial on the 2^nd^ extinction day there are no differences observed in low theta (**D**) or high theta (**E**) network responses, and during the final recall trial there are no differences observed in low theta (**F**) or high theta (**G**) network responses. In the FC, during the 1^st^ extinction trial on the 1^st^ extinction day there are no differences observed in low theta (**H**) or high theta (**I**) network responses, during the 1^st^ extinction trial on the 2^nd^ extinction day there are no differences observed in low theta (**J**) or high theta (**K**) network responses, and during the final recall trial there are no differences observed in low theta (**L**) or high theta (**M**) network responses.

**Supplemental Figure 4.**
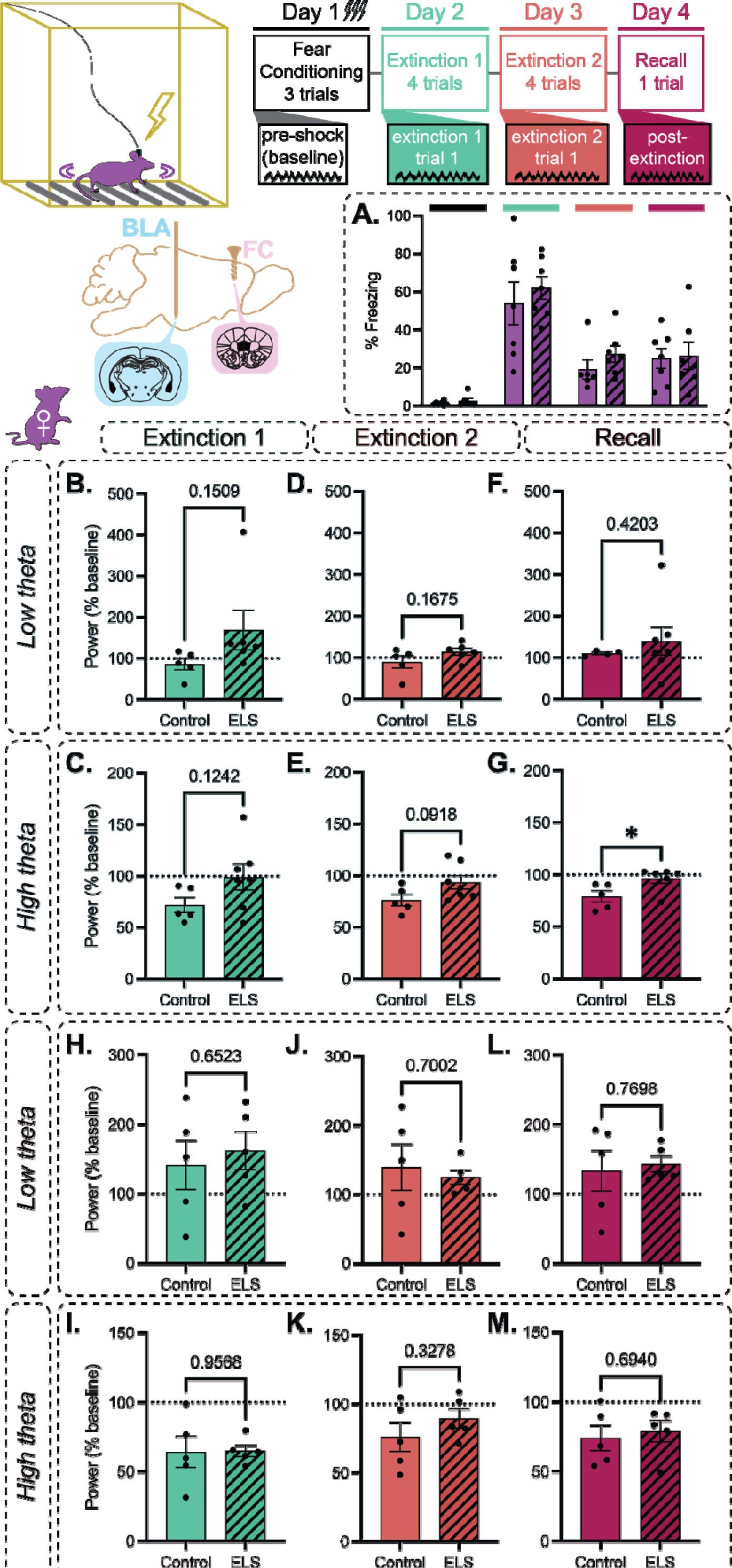
ELS alters BLA network processing during fear memory suppression in females. Female ELS mice are not significantly different from CTL in terms of freezing behavior across fear conditioning and extinction trials (**A**). In the BLA, during the 1^st^ extinction trial on the 1^st^ extinction day there are no differences observed in low theta (**B**) or high theta (**C**) network responses and during the 1^st^ extinction trial on the 2^nd^ extinction day there are no differences observed in low theta (**D**) or high theta (**E**) network responses. During the final recall trial, while no differences are observed in low theta (**F**), high theta network processing is significantly impacted (**G**). In the FC, during the 1^st^ extinction trial on the 1^st^ extinction day there are no differences observed in low theta (**H**) or high theta (**I**) network responses, during the 1^st^ extinction trial on the 2^nd^ extinction day there are no differences observed in low theta (**J**) or high theta (**K**) network responses, and during the final recall trial there are no differences observed in low theta (**L**) or high theta (**M**) network responses.

To investigate the impact of ELS on network processing of ethological threat, mice were single housed and exposed to predator odor (bobcat urine). CTL and ELS mice of both sex demonstrated a robust reduction in BLA LFP power across theta lasting ∼1 hr, however we focused on high theta because the effect was strongest within that range. Male mice demonstrated a fixed effect of time post exposure in the LFP in the BLA and FC, however there was no significant effect of stress condition in either brain region (Fig. 5). There were significant effects of both time post exposure and stress condition in the BLA LFP in females (Supp. Fig. 5A), with no difference observed in the FC LFP (Supp. Fig. 5B).

**Figure 5.**
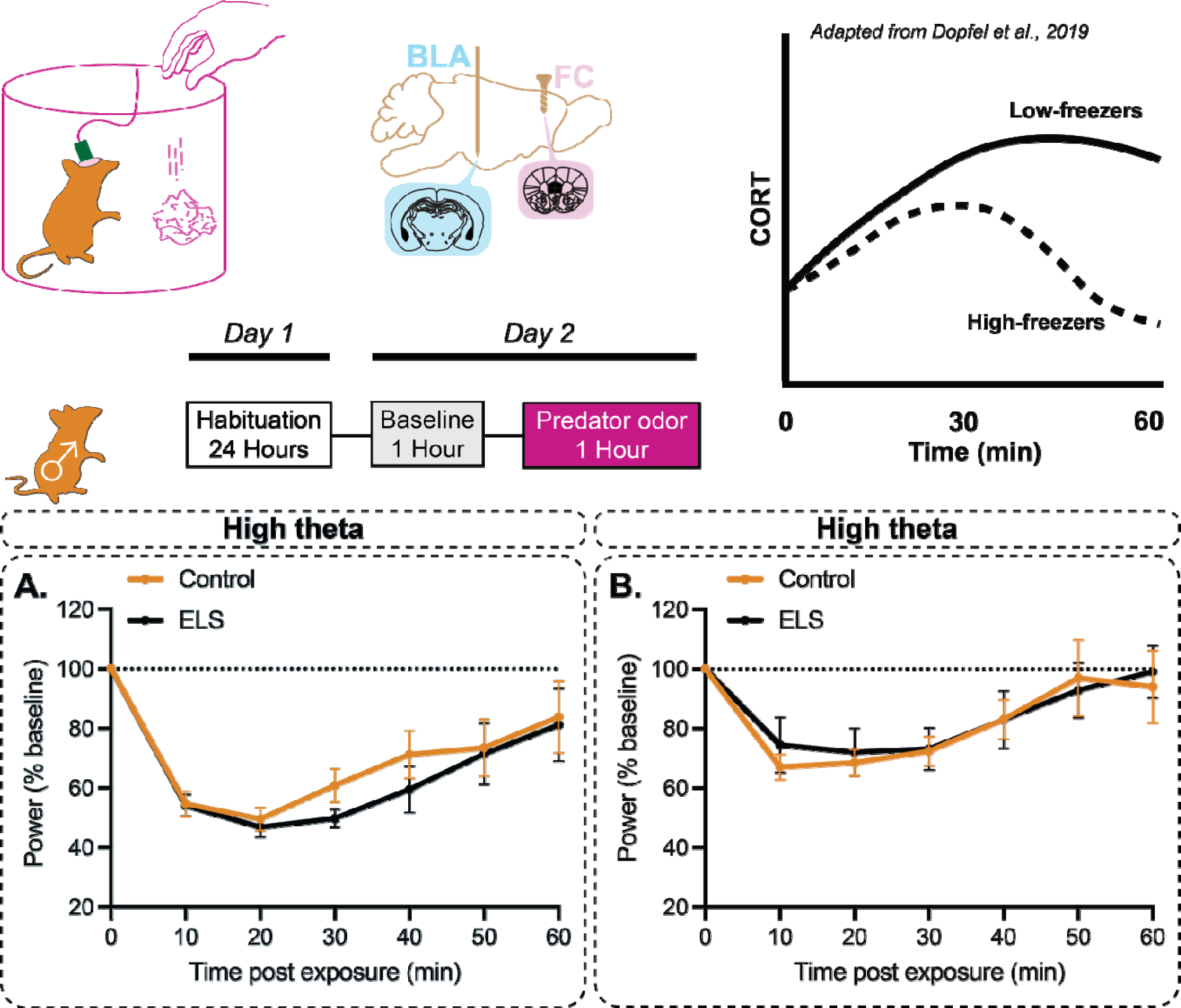
ELS does not impact network processing of predator odor in males. Male mice display significant suppression of BLA and FC high theta during exposure to predator odor on a timeline that is consistent with previous reports of post-odor CORT (Dopfel et al., 2019) (top right). ELS does not, however, impact the network response to predator odor in males in either the BLA (**A**) or the FC (**B**).

**Supplemental Figure 5.**
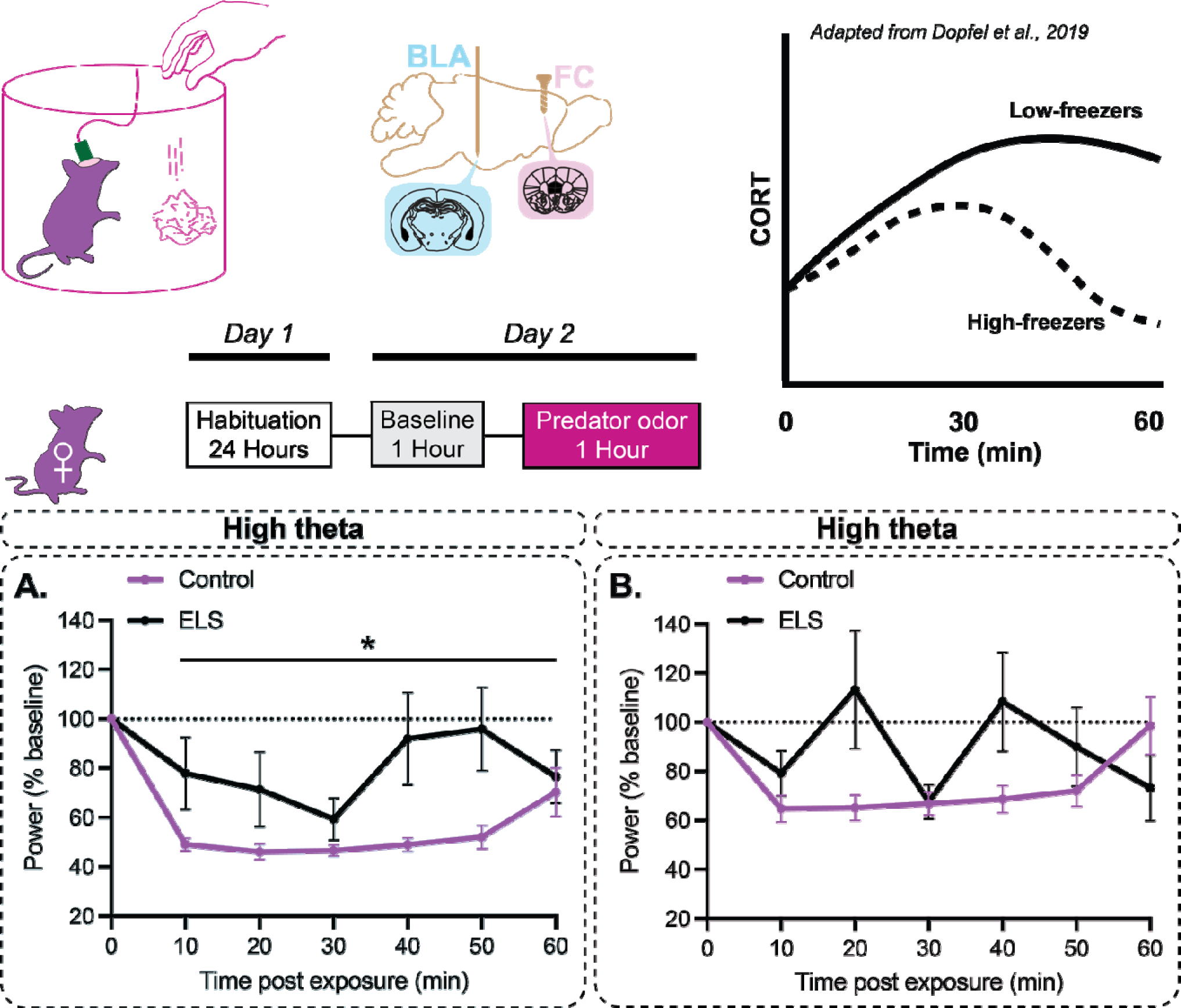
ELS alters network processing of predator odor in the BLA of females. Female mice display significant suppression of BLA high theta during exposure to predator odor on a timeline that is consistent with previous reports of post-odor CORT (Dopfel et al., 2019) (top right). ELS significantly impacts this effect in the BLA (**A**) but has no effect on post-odor high theta network responses in the FC (**B**) of females.

To investigate the impact of ELS on network processing of ethological threat contributing to altered behavior and survival, CTL and ELS mice were subjected to a looming disk apparatus in which a rapidly expanding black circle was triggered on a computer monitor placed over the cage. A small shelter was placed in the cage, which mice could dart to for shelter following stimulus onset. Other behavioral strategies observed included tail rattle and freezing. In most cases, when darting and tail rattle were observed they followed a brief (<1 sec) bout of freezing. For this reason, and to increase statistical power for less frequently observed behaviors, we grouped behaviors into active (darting and tail rattle) or inactive (prolonged freezing). During active responses, male ELS mice did not display any differences in BLA low theta state transitions (Fig. 6A), however ELS mice demonstrated a shift in high theta power which was significantly larger than CTL (Fig. 6B). No changes were observed at either frequency band in the FC (Fig. 6C-D). During inactive coping, male CTL mice displayed a reduction in low theta power which was significantly different from ELS mice (Fig. 6E), while state shifts in high theta were not significantly different (Fig. 6F). Like with active responses, no significant effects were observed in the FC (Fig. 6G-H). In females, active responses were not associated with significant state shifts across low or high theta in the BLA or FC (Supp. Fig. 6 A-D). Inactive responses, however, were associated with significantly different network state transitions in ELS mice in BLA low theta (Supp. Fig. 6E), but not high theta (Supp. Fig. 6F). Conversely, in the FC, low theta transitions were not significantly different in low theta (Supp. Fig. 6G), but were significantly different in high theta (Supp. Fig. 6H).

**Figure 6.**
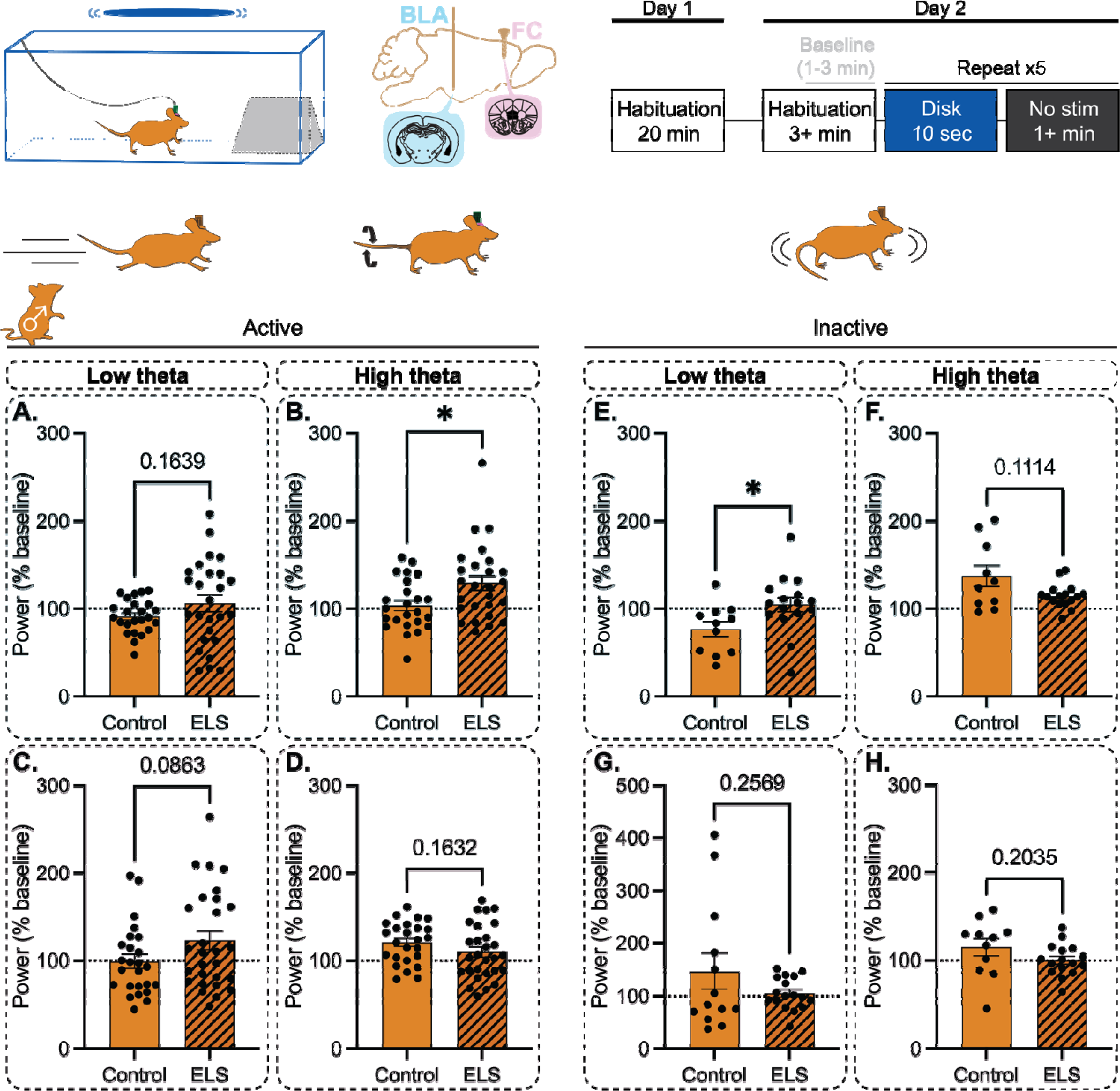
ELS significantly alters network responses during active and inactive behavioral strategies for responding to a looming disk in males. Active behaviors include darting and rattling while freezing is referred to as inactive behavior. In males, ELS does not impact low theta network responses in the BLA during active behaviors (**A**), however BLA high theta responses are significantly different (**B**). No differences are reported in FC low theta (**C**) or high theta (**D**). During inactive behavior, males display significantly different BLA low theta network responses (**E**), with no changes in high theta (**F**). In the FC, there are no differences in low theta (**G**) or high theta (**H**).

**Supplemental Figure 6.**
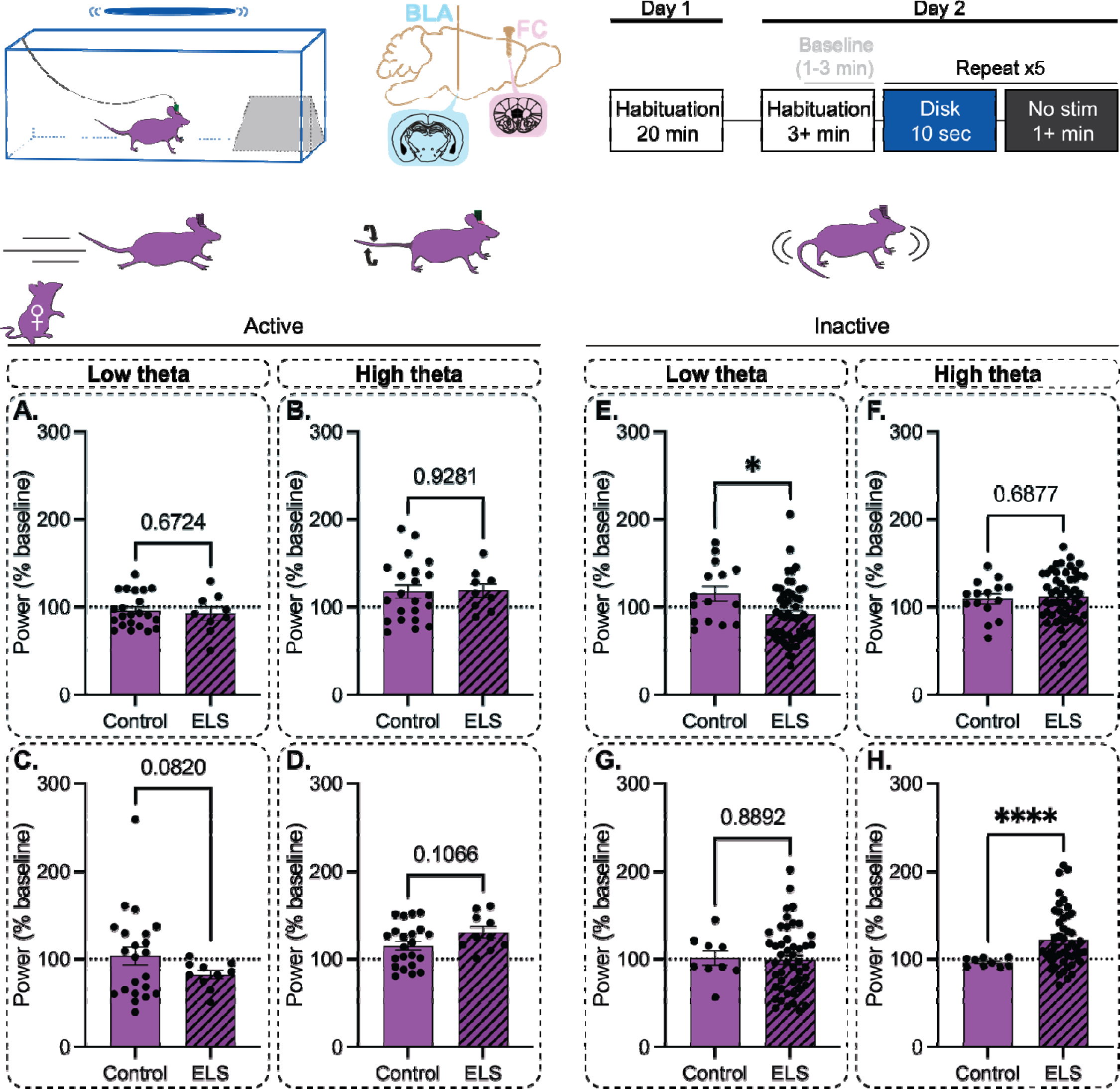
ELS significantly alters network responses during inactive behavioral strategies for responding to a looming disk in females. Active behaviors include darting and rattling while freezing is referred to as inactive behavior. In females, ELS does not impact network responses in the BLA during active behaviors in the range of low theta (**A**) or high theta (**B**). Similarly, no differences are reported in FC low theta (**C**) or high theta (**D**). During inactive behavior, females display significantly different BLA low theta network responses (**E**), with no changes in high theta (**F**). In the FC, females display no changes in low theta (**G**), however there is a significant difference in high theta network responses (**H**).

Previous studies have implicated changes in BLA PV interneuron expression during development in mediating the adverse effects of developmental adversity (Manzano Nieves et al., 2020). Furthermore, our lab has demonstrated that these interneurons are involved in processing high stress information (Davis et al., 2017) and are capable of driving BLA network states (Antonoudiou et al., 2022). PNNs are developmentally regulated and have been described as preferentially targeting PV interneurons. For these reasons, we decided to investigate whether ELS impacts developmental BLA PV and or PNN expression in males or females (Fig. 7A). We did not observe any differences between CTL and ELS mice in terms of PV density, PNN density, or percentage of PV interneurons coexpressing PNNs in males (Fig. 7B-D) or females (Fig. 7E-G).

**Figure 7.**
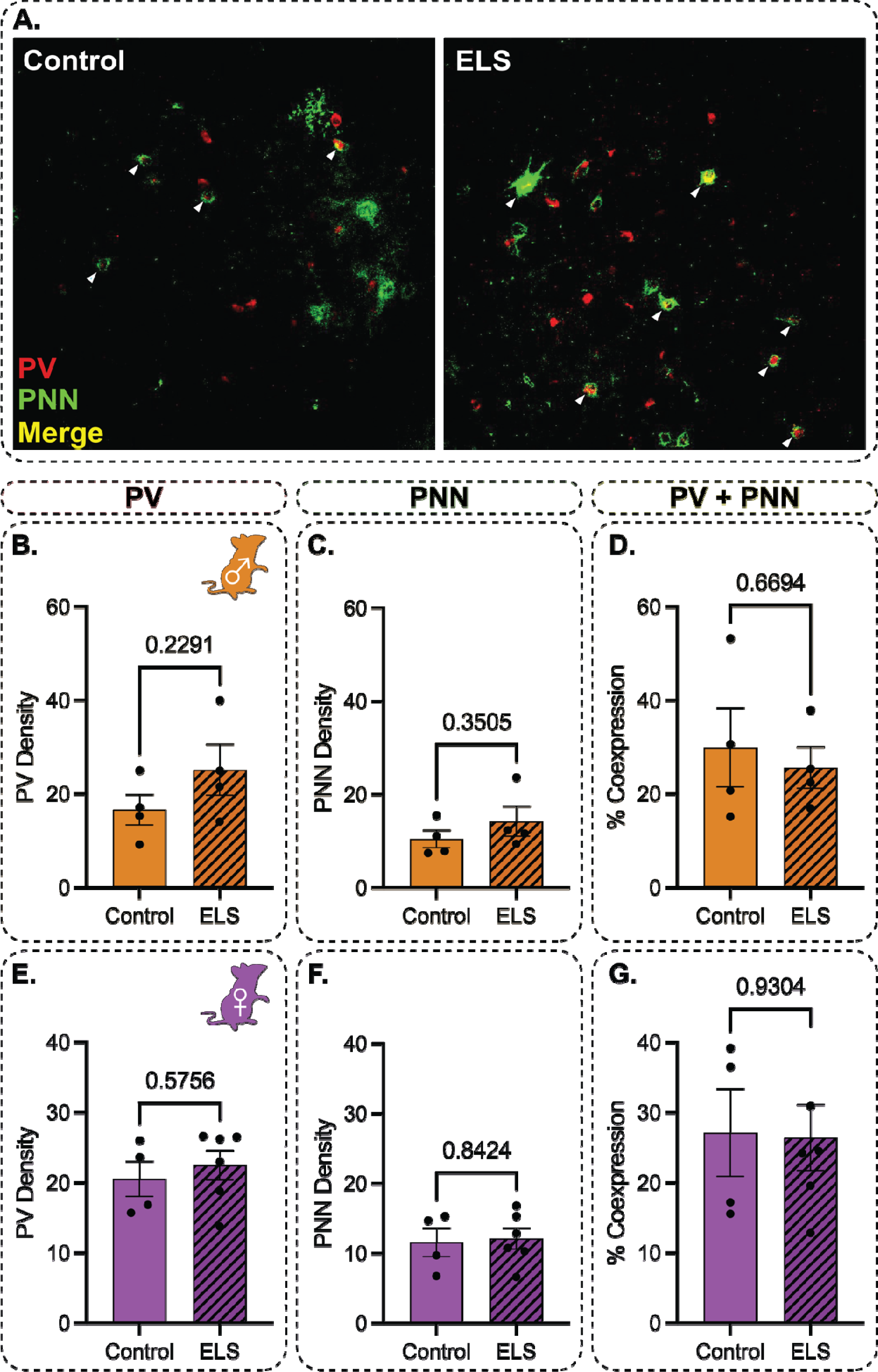
ELS does not impact developmental PV expression or PNN targeting in males or females at 4 weeks postnatal. Representative images of PV and PNN histology (**A**). In males, ELS does not significantly impact PV expression (**B**), PNNs (**C**), or PNN targeting of PV interneurons (**D**). Similarly, in females, ELS does not significantly impact PV expression (**B**), PNNs (**C**), or PNN targeting of PV interneurons (**D**).

To look for functional changes in cellular physiology contributing to behavioral and network effects of ELS, we used whole cell patch clamp in the current clamp configuration to record activity from principal and PV neurons in the BLA. To confirm that we were targeting separate populations, we pooled data across sex and stress condition for our initial analyses. BLA principal and PV neurons were significantly different from one another in terms of input/output slope (Supp. Fig. 7A-B), peak firing rate (Supp. Fig. 7C), half-width (Supp. Fig. 7D), and action potential peak (Supp. Fig. 7E). We also observed that PV interneurons are significantly more responsive to input during low frequency membrane oscillations than principal neurons (Supp Fig. 7F-G), however there were no differences during gamma oscillations (Supp Fig. 7H).

**Supplemental Figure 7.**
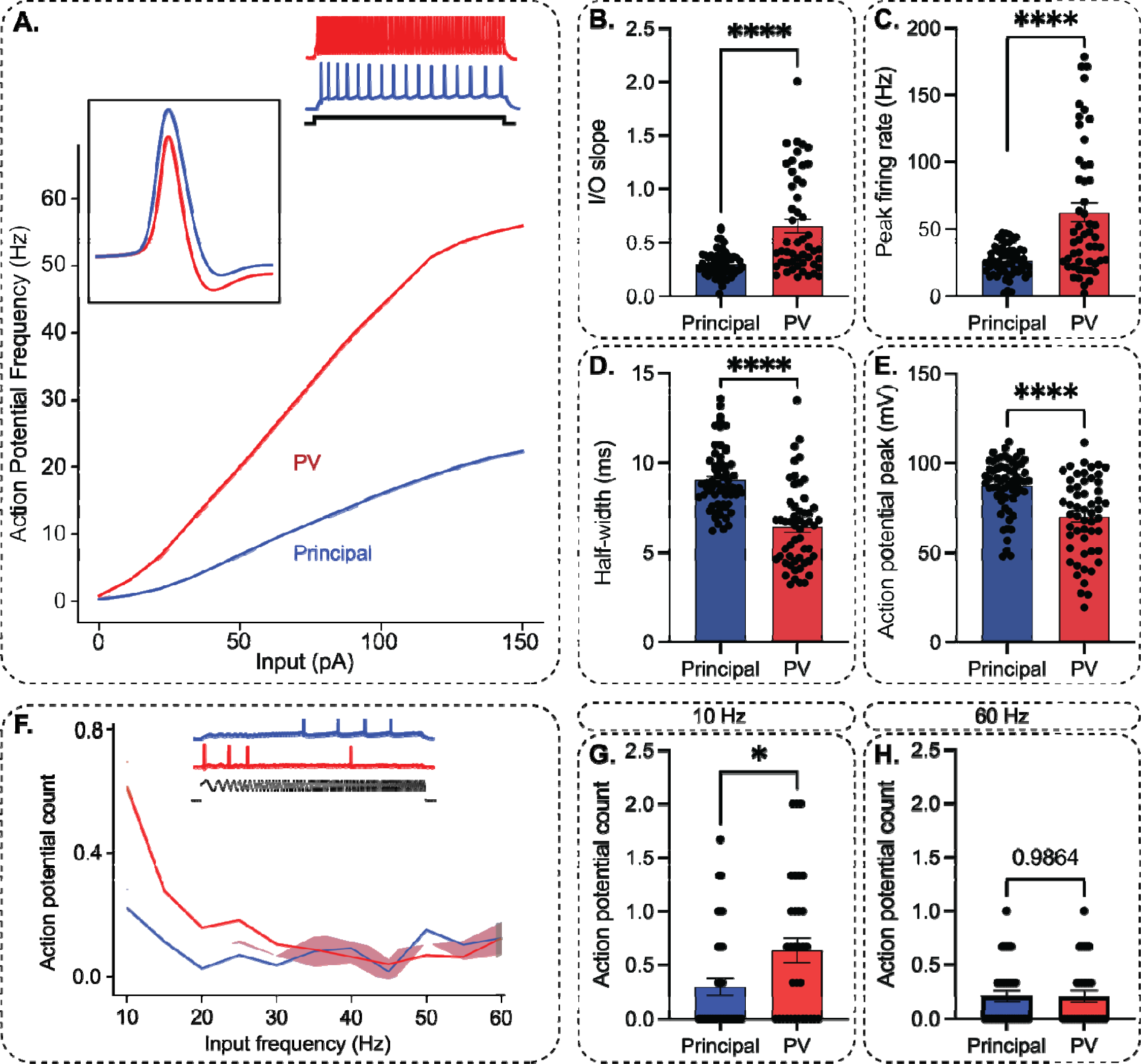
BLA PV interneurons and principal neurons segregate into discrete populations with distinct resonance profiles. Averaged I/O plots (**A**). PV interneurons in the BLA display significantly increased input/output slope (**B**) and peak firing rate (**C**), as well as significantly reduced half width (**D**) and action potential peak (**E**) as compared to principal neurons. Averaged resonance profiles (**F**) reveal that PV interneurons are significantly more active than principal neurons during suprathreshold theta stimulation (**G**), while there are no differences in recruitment during gamma stimulation (**H**).

When separated by sex, male ELS mice demonstrated no differences in BLA PV interneuron input/output slope (Fig. 8A-B), peak firing rate (Fig. 8C), afterhyperpolarization (AHP) (Fig. 8D), or action potential peak (Fig. 8E). Male ELS mice were also not significantly different with regard to principal neuron input/output slope (Fig. 8F-G), however ELS mice demonstrated a significantly reduced peak firing rate (Fig. 8H). No differences were observed in AHP (Fig. 8I) or action potential peak (Fig. 8J). Female mice did not display any significant differences in BLA PV interneuron input/output slope (Supp. Fig. 8A-B), peak firing rate (Supp. Fig. 8C), AHP (Supp. Fig. 8D), or action potential peak (Supp. Fig. 8E). Similarly, no differences were observed in female principal neuron input/output slope (Supp. Fig. 8G), peak firing rate (Supp. Fig. 8H), AHP (Supp. Fig. 8I), or action potential peak (Supp. Fig. 8J).

**Figure 8.**
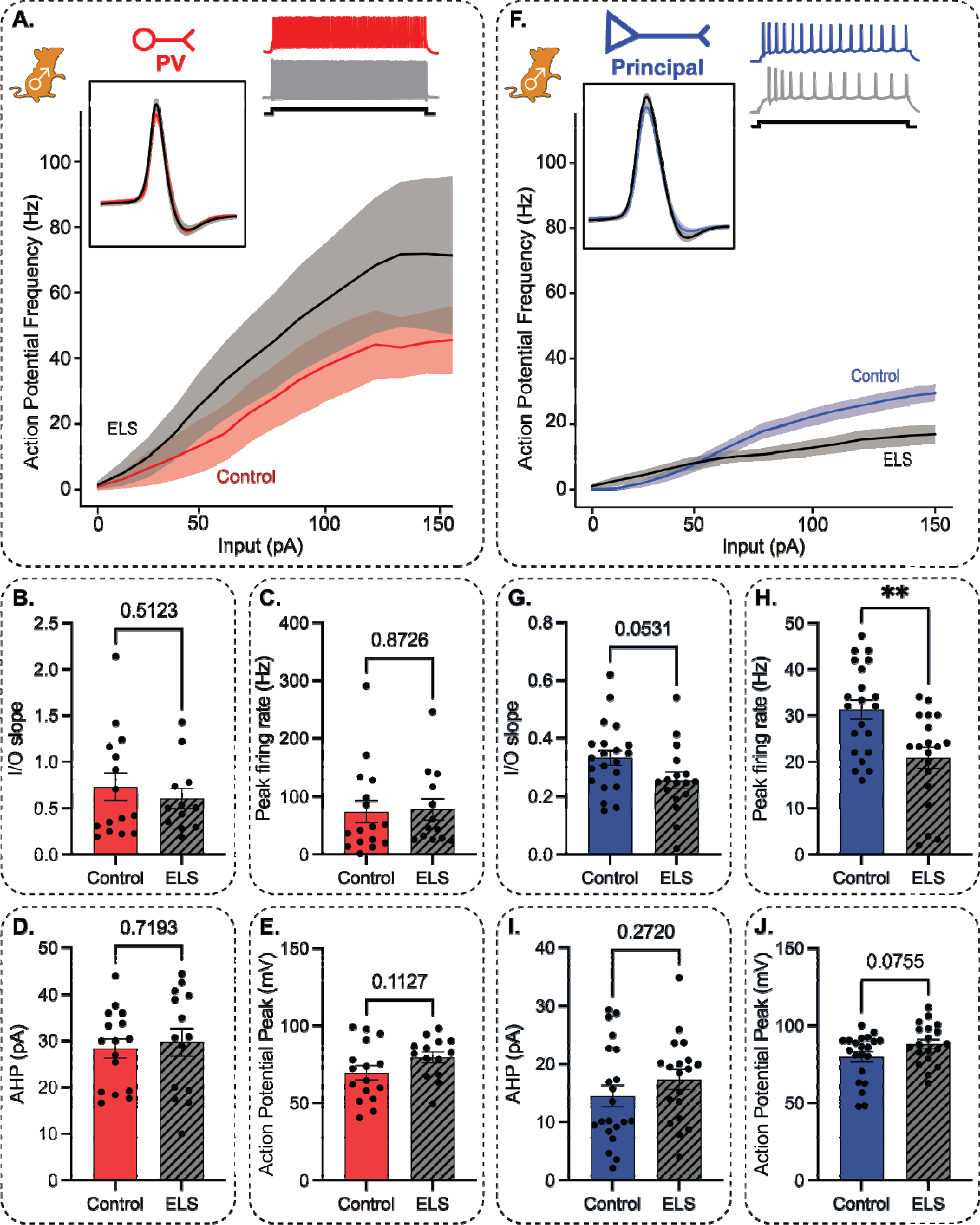
Effects of ELS in males are mediated by altered BLA principal neuron active properties. Representative input/output curves (main), waveforms (top left), and response following membrane depolarization (top right) for male BLA PV interneurons (**A**) and principal neurons (**F**). Male ELS PV interneurons are not significantly different from CTL in terms of input/output slope (**B**), peak firing rate (**C**), afterhyperpolarization (**D**), or action potential peak (**E**). Male ELS principal neurons are also not significantly different in terms of input/output slope (**G**) but display a significantly reduced peak firing rate compared to CTL (H). No differences are reported regarding afterhyperpolarization (**I**) or action potential peak (**J**).

**Supplemental Figure 8.**
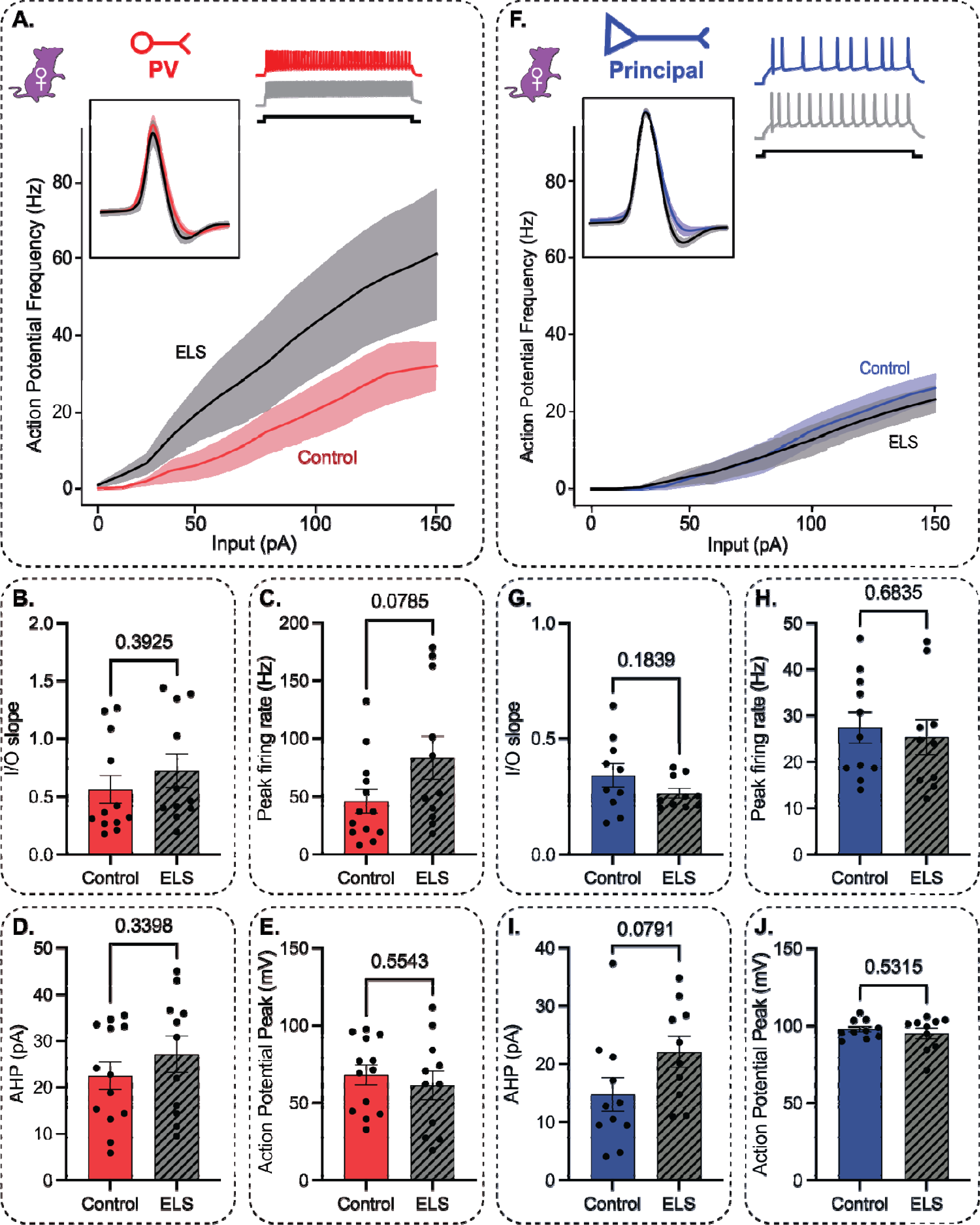
Effects of ELS in females do not result from changes in BLA PV interneuron or principal neuron activation properties measured using standard current steps. Representative input/output curves (main), waveforms (top left), and response following membrane depolarization (top right) for female BLA PV interneurons (**A**) and principal neurons (**F**). Female ELS PV interneurons are not significantly different from CTL in terms of input/output slope (**B**), peak firing rate (**C**), afterhyperpolarization (**D**), or action potential peak (**E**). Similarly, female ELS principal neurons are also not significantly different from CTL in terms of input/output slope (**G**), peak firing rate, (**H**), afterhyperpolarization (**I**) or action potential peak (**J**).

When presented with a suprathreshold chirp stimulus, PV cells from male ELS mice were not different from CTL in terms of action potential generation at low frequencies (Fig. 9B) or high frequencies (Fig. 9C). Principal neurons from male ELS mice, however, were significantly more active than CTL during low frequency membrane oscillations (Fig. 9E) but displayed no differences during high frequency oscillations (Fig. 9F). Conversely, PV cells from female mice were significantly more active than CTL during low frequency oscillations (Supp. Fig. 9B) and were significantly less active during high frequency oscillations (Supp. Fig. 9C). Principal neurons from female ELS mice were not significantly different from CTL in activity either during low frequency oscillations (Supp. Fig. 9E) or high frequency oscillations (Supp. Fig. 9F). Collectively, these data demonstrate functional changes in the cellular physiology of neurons in the BLA which corresponds to altered oscillatory states associated with distinct behavioral outcomes following ELS in males and females.

**Figure 9.**
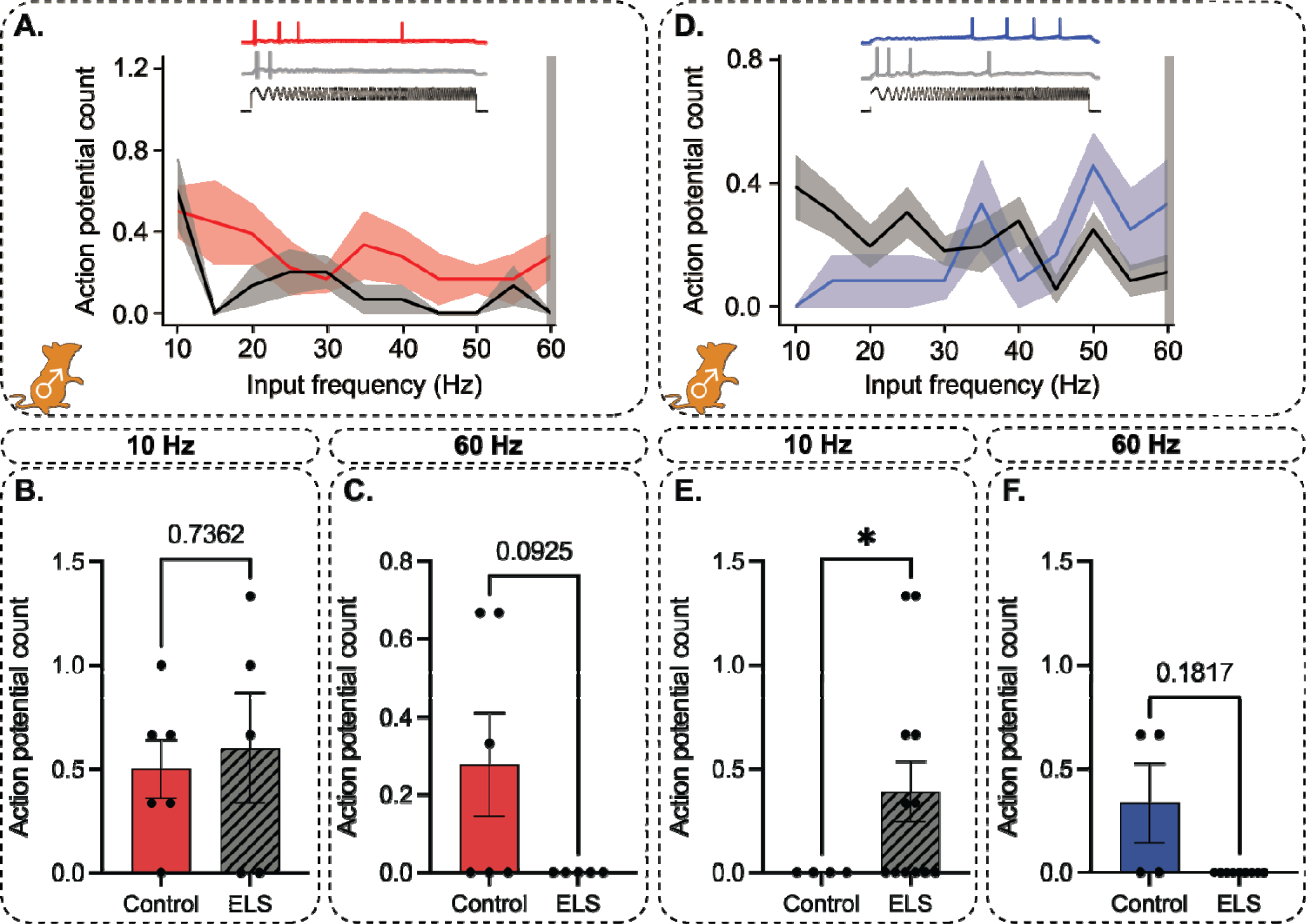
Impacts of ELS in males are mediated by altered BLA principal neuron resonance profiles. Representative resonance profiles across stimulation frequencies (main) and representative voltage traces from CTL (top-top) and ELS (top-middle) mice following a suprathreshold chirp stimulation (top-bottom) in male BLA PV interneurons (**A**) and principal neurons (**D**). ELS PV interneurons are not significantly different from CTL in terms of action potential generation during low frequency (**B**) or high frequency (**C**) stimulation. Male ELS principal neurons are significantly more active than CTL during low frequency stimulation (**E**) but are not differentially active during high frequency stimulation (**F**).

**Supplemental Figure 9.**
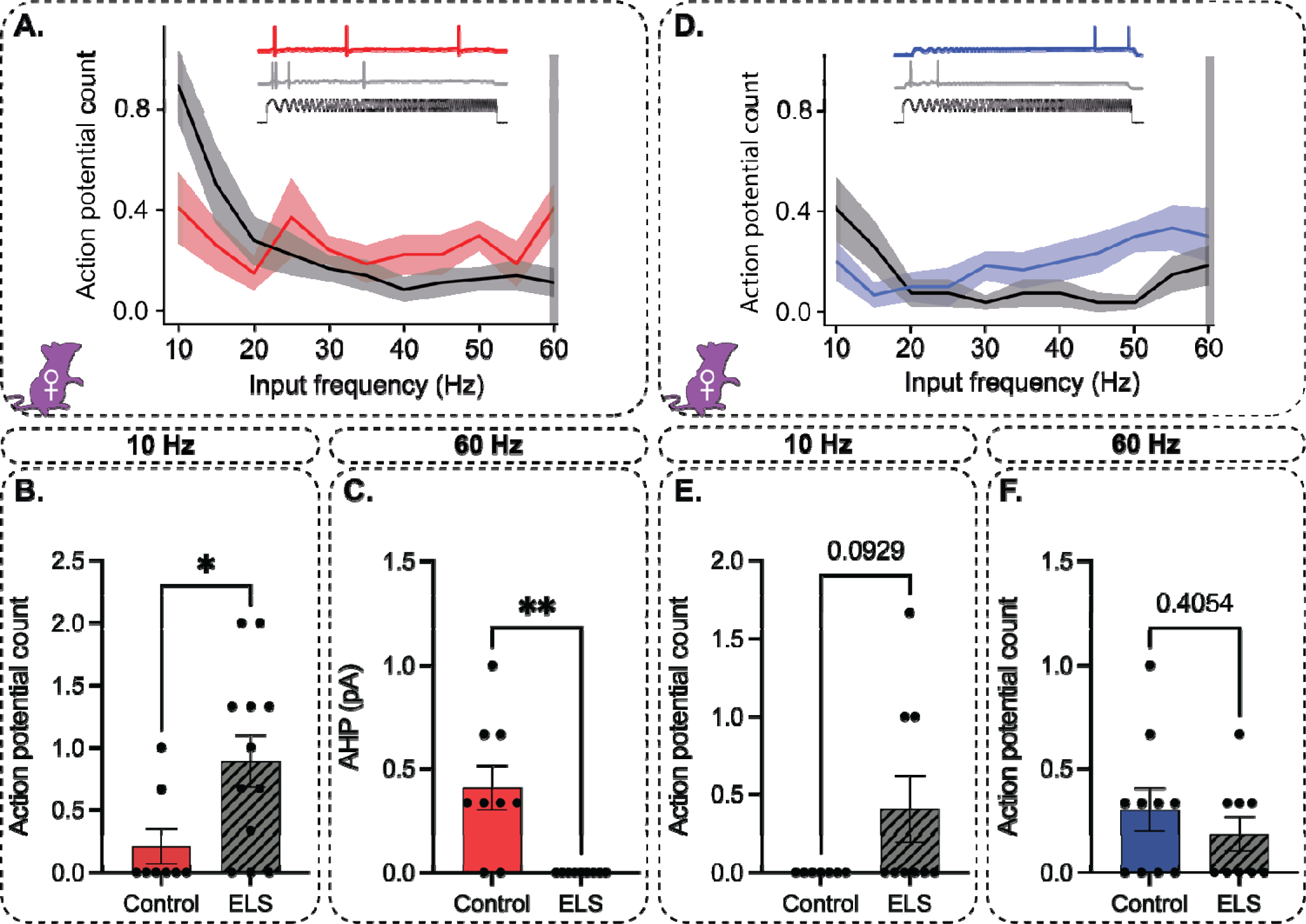
Impacts of ELS in females are mediated by altered PV interneuron resonance profiles. Representative resonance profiles across stimulation frequencies (main) and representative voltage traces from CTL (top-top) and ELS (top-middle) mice following a suprathreshold chirp stimulation (top-bottom) in female BLA PV interneurons (**A**) and principal neurons (**D**). Female ELS PV interneurons are significantly more active during low frequency stimulation (**B**) and less active during high frequency stimulation (**C**) when compared to CTL. Conversely, female ELS principal neurons are not significantly different in terms of activity during low frequency (**E**) or high frequency (**F**) stimulation.

## Discussion

Here we demonstrate that a murine MS ELS model results in sex-specific changes to local network theta states across relevant brain regions during critical processing of stressful information. Changes to these network states have previously been associated with altered behavioral strategies for coping with stress (Antonoudiou et al., 2022) and conditioned fear (Davis et al., 2017), however the contribution of adolescent trauma has not been explored. We further suggest that these changes result from altered BLA projection neuron and interneuron neuron physiology in males and females, respectively.

We report that ELS results in changes to behavioral strategies for escaping high stress environments in males, both during adolescence and adulthood (Fig. 2A-D). Concurrently, male ELS mice display a diminished stress response following acute restraint in adulthood (Fig. 2E). This relationship between CORT and immobility in the FST is consistent with previous reports (Fenton et al., 2015; X. Xie et al., 2020). Interestingly, female ELS mice do not appear to be impacted in the context of these behavioral paradigms or neuroendocrine deficits (Supp. Fig. 1). To determine genetic contributions to the behavioral effects of ELS we raised DBA2J mice under identical conditions. ELS mice were not different in terms of percentage of time spent in the center vs. periphery of the open field in either males (Supp. Fig. 2A) or females (Supp. Fig. 2B). Similarly, no differences were observed in percentage of total time spent with the resident in the SI test in either males (Supp. Fig 2C) or females (Supp Fig. 2D). However, when the impacts of stress and strain were compared in males, 2-way ANOVA reveals significant differences between DBA/2J and C57/BL6J ELS mice, suggesting that genetic background may contribute to stress vulnerability and resilience.

Next, we demonstrate that male BLA and FC LFP theta power is not significantly impacted by ELS under non-stressful conditions (Fig. 3). The ratio of low-to-high theta has previously been used to describe network states contributing to internal anxiety states (Davis et al., 2017), so we included this metric in our analysis. Females display similar patterns in the BLA (Supp. Fig. 3A-D), however ELS reduces high theta and increases the low-to-high theta ratio (Supp. Fig 3E-H). Collectively, these data suggest that ELS may contribute to altered network states guiding internal anxiety states at rest in females, but not males. Interestingly, it has been shown that repetitive transcranial magnetic stimulation (rTMS) – which is used in humans to alter brain circuitry at rest and treat depression – may be more effective in females than males (Desai et al., 2023).

We further demonstrate that ELS does not impact freezing patterns or network state transitions in the BLA or FC of males following contextual fear conditioning (Fig. 4). In the case of females, while freezing behavior is unaffected (Supp. Fig. 4A), CTL mice demonstrate a change in BLA high theta oscillations (relative to a pre-shock trial) during recall which is significantly lower than the change seen in ELS mice (Supp. Fig. 4G). While freezing is not significantly different during this timepoint, it is possible that the altered network states reflect a primed state to respond to threat exposure although the behavioral output is similar.

We additionally show that exposure to predator odor drives a robust decrease in high theta oscillations in the BLA and FC lasting ∼1 hr. In males, ELS does not impact this reduction in power in either region (Fig 5). In females, however, ELS significantly dampens this response in the BLA (Supp. Fig. 5A). These results suggest that females may be more vulnerable to network changes following prolonged exposure to ethological threat.

In response to an acute predator stimulus with multiple behavioral survival strategies, we report that male ELS mice display increased high theta power over baseline in the BLA relative to CTLs during active responses (Fig. 6B). During inactive responses however, the BLA low theta response is significantly altered (Fig. 6E). Female CTL and ELS mice demonstrate no differences in low or high theta state shifts during active responding in the BLA or FC (Supp. Fig. 6A-D), however inactive responding corresponds with significantly altered state shifts across low theta in the BLA (Supp. Fig. 6E) and high theta in the FC (Supp. Fig. 6H). These data are consistent with and further our knowledge regarding the role of FC and BLA network states in guiding behavioral responses to stressful stimuli.

Previous studies have implicated altered BLA PV interneuron expression as a mediator of the effects from a different model of ELS (Manzano Nieves et al., 2020). Here, we demonstrate that effects of a MS paradigm on behavioral and network states occur independent of changes in BLA PV expression at a 4 week timepoint (Fig. 7). To address cell-specific changes in terms of physiology and implications for circuit engagement, we recorded from PV and principal BLA neurons. While PV interneuron physiology remains unaffected by ELS in males (Fig. 8A-E), principal neurons display reduced peak firing following depolarization (Fig. 8H) suggesting suppression of a specific output circuit although it is also possible that the changes may be more general. Interestingly, ELS also increases recruitment of principal neurons during theta-range membrane oscillations in males (Fig 9E). In females, ELS does not impact PV or principal neuron physiology in response to square pulse current steps (Supp. Fig. 8), however ELS PV interneurons were significantly more responsive to theta membrane oscillations (Supp. Fig. 9B) and less responsive to gamma oscillations (Supp. Fig. 9C). These findings demonstrate sex-dependent impacts of ELS on BLA cellular physiology and may contribute to the divergent network and behavioral changes observed following ELS in males vs. females.

Collectively, our findings provide critical insights regarding the influence of developmental adversity on network and behavioral states. Changes in dynamic BLA-FC network states are increasingly recognized as being important mediators of psychiatric vulnerability, however this is the first description of the impact of ELS on BLA and FC theta oscillations during acute ethological stress and conditioned fear. Our findings reveal strong sex differences in the cellular, network, and behavioral consequences of developmental adversity, suggesting that perturbations in BLA and FC network states and circuit output may contribute to the pathophysiology and targeted manipulations of these networks may be effective at treating stress related psychiatric conditions under specific conditions.

## Acknowledgements

The authors would like to thank Drs. Kevin Bath, John Christianson, and Heidi Meyer for providing feedback on experimental design. The authors would also like to thank Rachel Rivas, Alice Fang, and Jenah Gabby for their contributions. This research was supported by the National Institute of Health under the following award numbers: F31MH133280, R01AA026256, R01NS105628, R01NS102937, R01MH128235, P50MH122379.

---

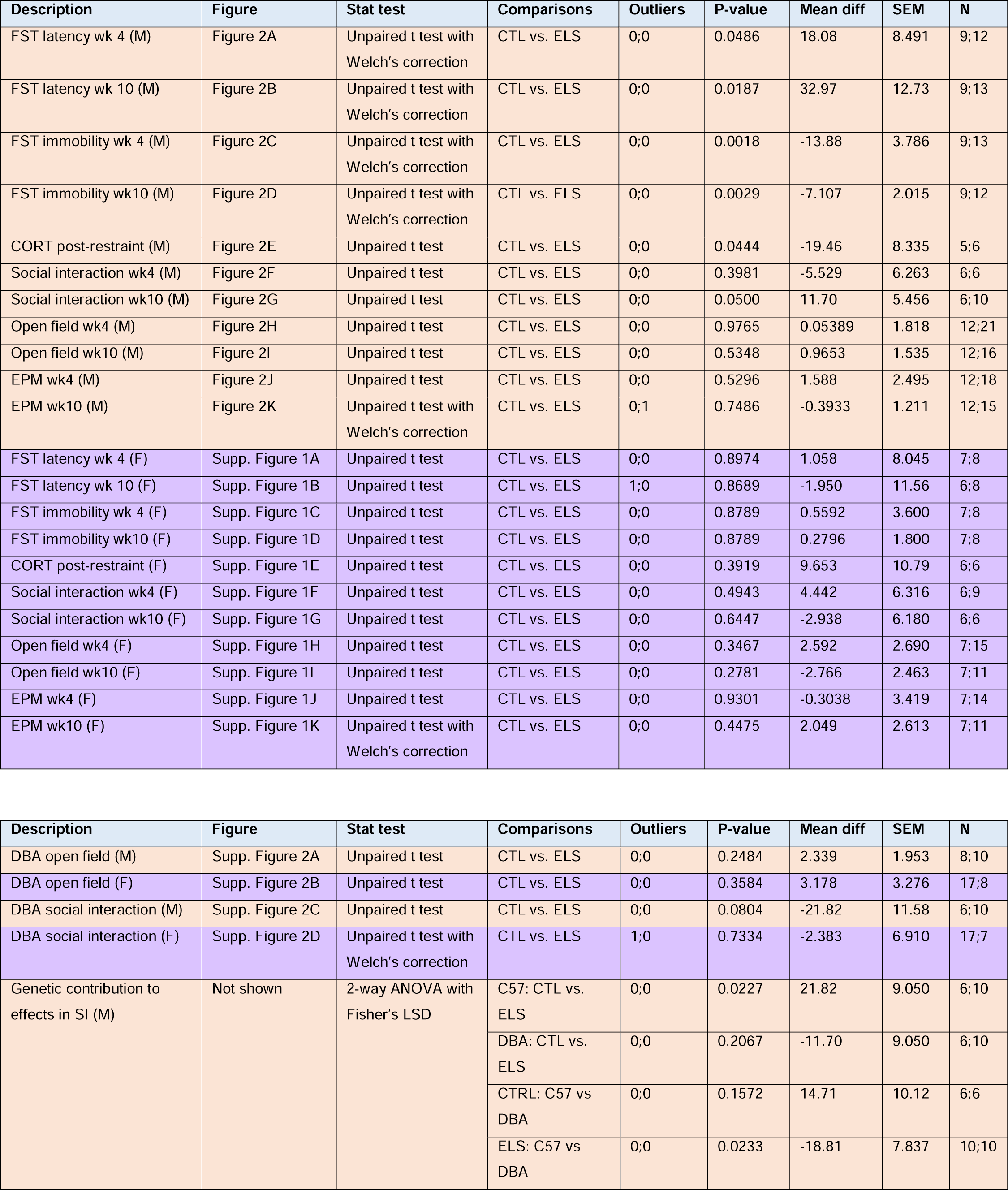
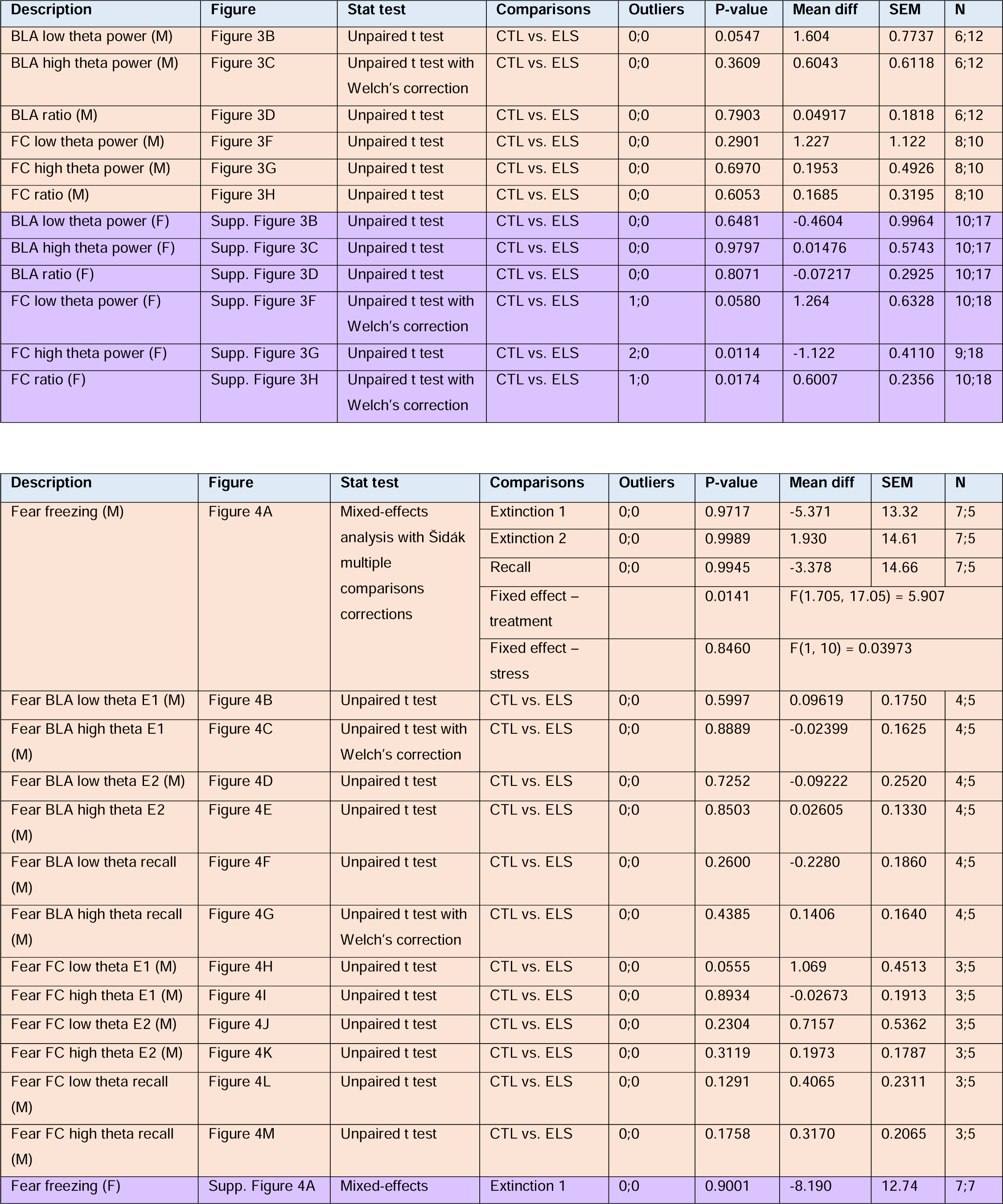
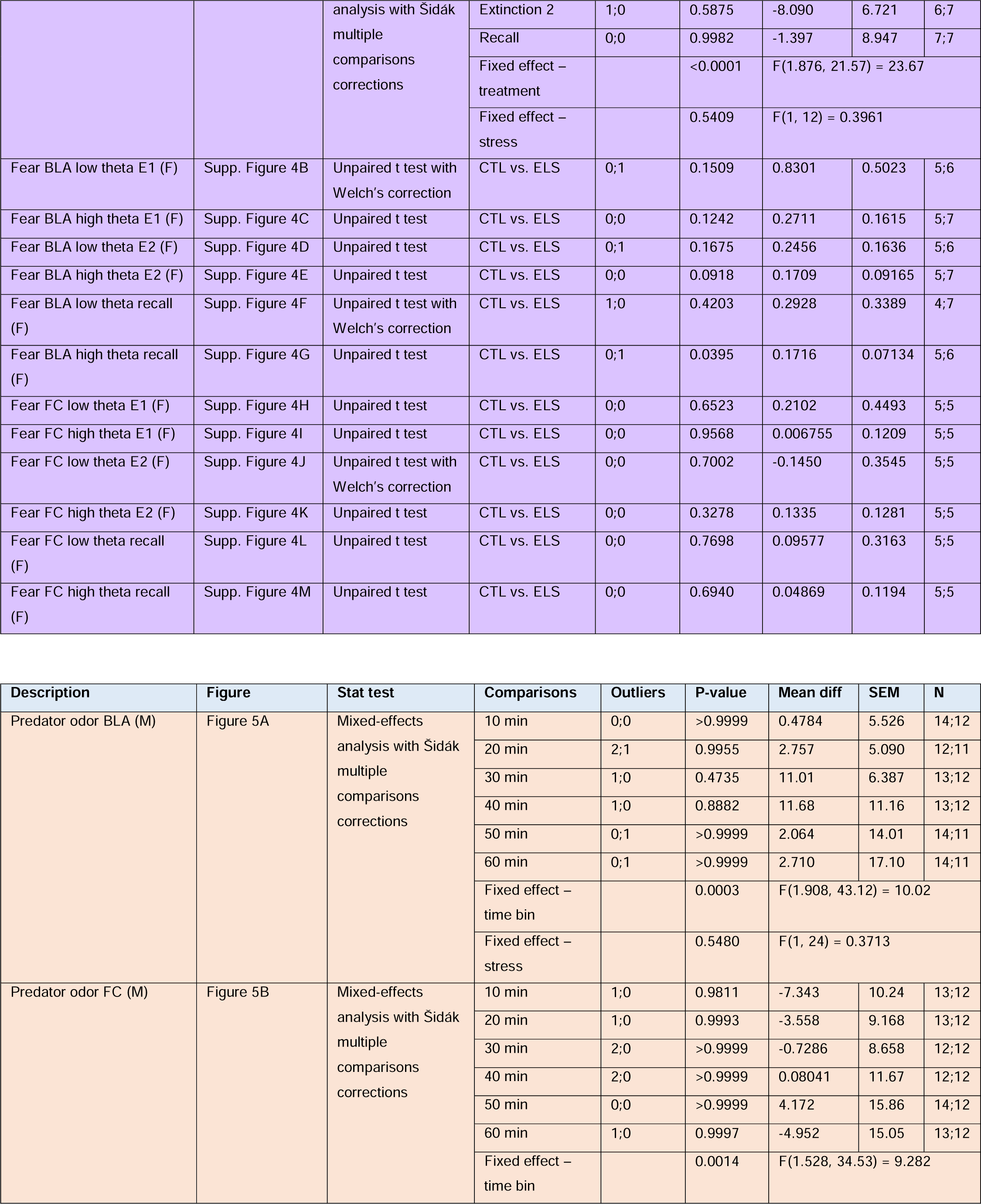
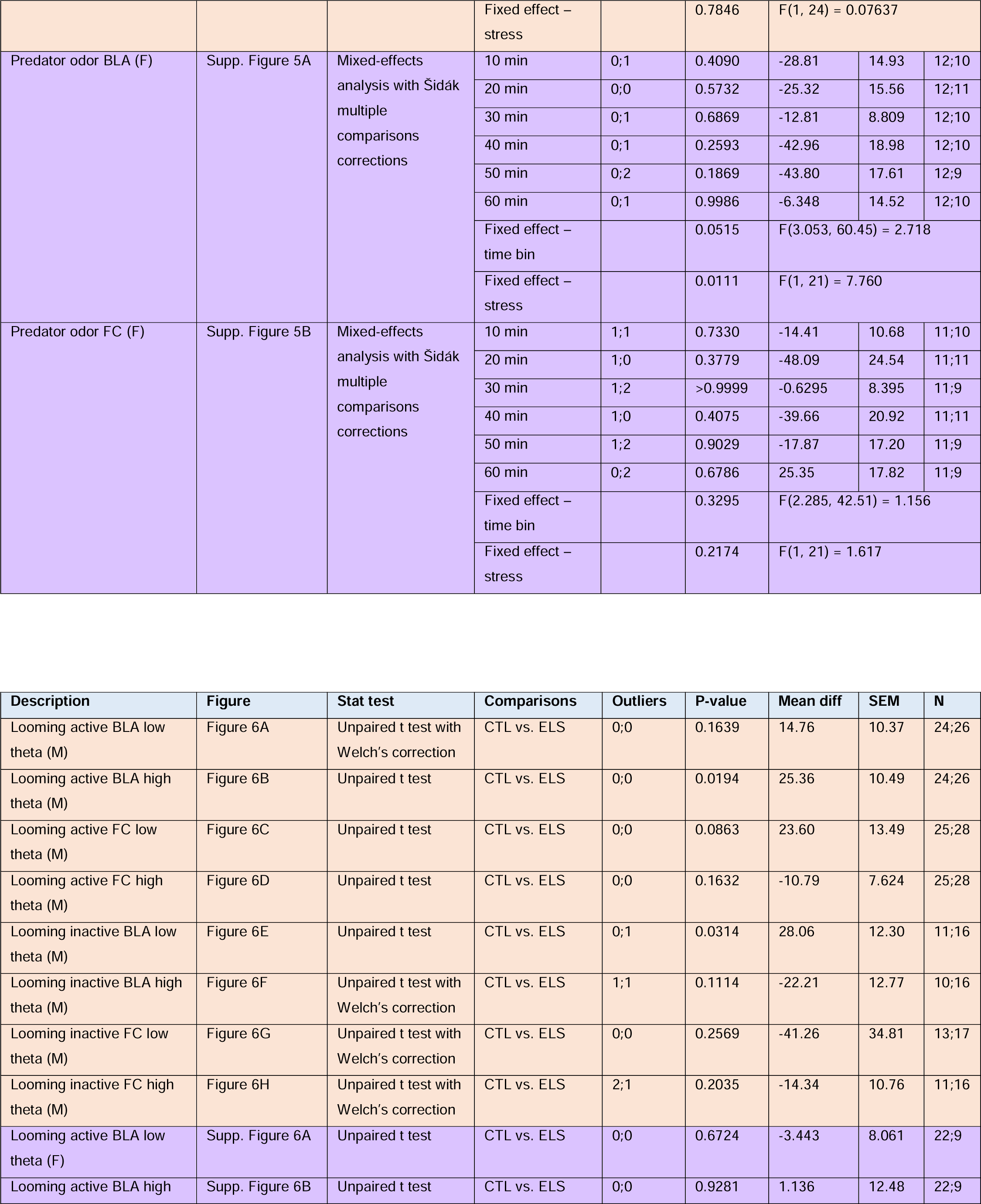
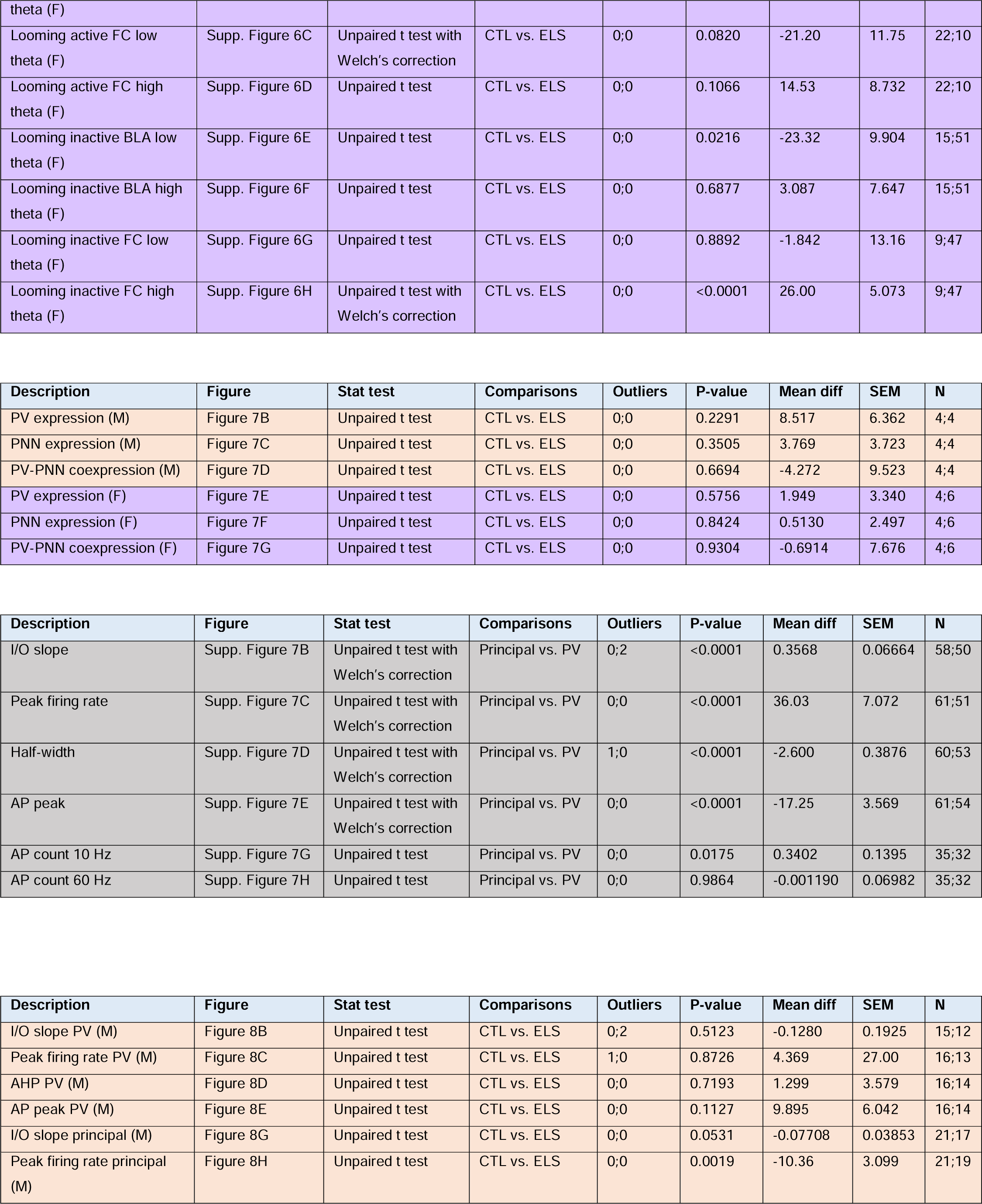
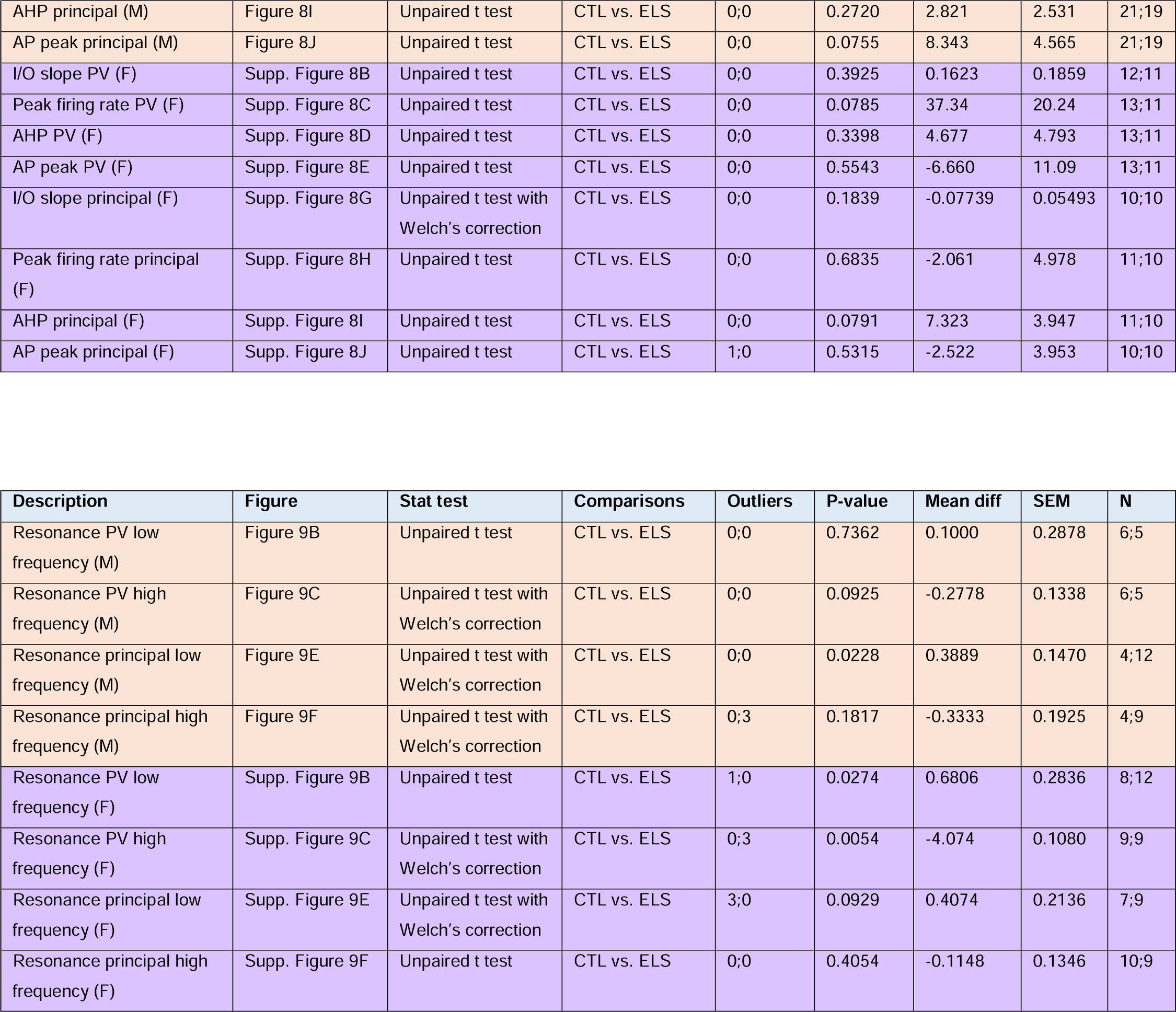

